# Integrated multi-omics reveals minor spliceosome inhibition causes molecular stalling and developmental delay of the mouse forelimb

**DOI:** 10.1101/2022.11.10.516037

**Authors:** Kyle D. Drake, Saren M. Springer, Kevon O. Afriyie, Tomas D. Lopes, Kaitlin N. Girardini, Rahul N. Kanadia

## Abstract

Developmental insults causing limb progenitor cell cycle defects or death tend to produce micromelic limbs with maintained segmentation. This suggests that the developing limb is plastic yet has a bias towards proximo-distal patterning. Here we use a minor spliceosome-deficient (U11-null) mouse forelimb, which has severe micromelia yet maintains proximo-distal segmentation, to decipher the mechanism(s) underlying this form of developmental robustness. We show that U11 loss triggers transcriptomic stalling upon spatially heterogenous mis-splicing of minor intron-containing genes. Through spatial transcriptomics, we detected a failure of the U11-null forelimb to separate its distal patterning program from its proximal differentiation program, which was supported by single-cell RNAseq-determined developmental delay of U11-null chondroprogenitors. Ultimately, these molecular and cellular deficits culminated in perturbed chondrogenesis, myogenesis, and axonogenesis. Taken together, we suggest that, upon sensing depletion of progenitors, the limb halts its transcriptional networks to pause its cellular trajectory, affording time to restructure its developmental program.

## Introduction

The limb is a biological tool designed for environmental interaction. Thus, species with distinct ecological niches tend to have diverse limb shapes and sizes^1,2^. These alterations operate within a fundamental, tripartite framework, which includes a stylopod (upper limb), zeugopod (middle limb), and autopod (distal limb)^1,2^. This observation suggests that the developing limb is plastic yet has a developmental bias favored towards its proximo-distal (shoulder to fingertip) segmentation. Evidence supporting this idea has been observed across a range of experimental models that induce limb progenitor cell (LPC) cell cycle defects and/or cell death^3–8^, including Wolpert’s classical X-irradiation experiment^9^ that was mechanistically resolved much later^10^. However, an integrated understanding of limb plasticity at the molecular and cellular level upon changes in LPC number remains unclear.

Recently, we reported that minor spliceosome inhibition in the developing mouse limb results in severe micromelia due to mis-splicing of minor intron containing genes (MIGs) that regulate cell cycle, thereby causing reduced proliferation and apoptosis of LPCs^11^. Despite these cellular deficits, proximo-distal segmentation was maintained^11^. This finding is in line with limb reduction defects that do not perturb limb patterning in multiple developmental disorders caused by disrupted minor spliceosome function, including microcephalic osteodysplastic primordial dwarfism type 1 (MOPD1), Roifman syndrome (RS), and Lowry-Wood syndrome (LWS)^12–16^. Thus, we reasoned that resolving how the minor spliceosome-deficient limb responds to loss of LPCs at a molecular, cellular, and systems level could lend insight toward the mechanism(s) underlying the plasticity and proximo-distal developmental bias of the limb.

Here, we conduct spatiotemporal multi-omics on the minor spliceosome-deficient (*Rnu11*^flx/flx^::*Prrx1-Cre*^+^) mouse forelimb^11,17^. Through temporal RNAseq, we report that the U11-null (mutant) forelimb shows sustained transcription of key limb patterning genes that are downregulated across time in the wild-type (WT) forelimb, which we define as transcriptomic stalling. Subsequently, we designed a novel microdomain analysis that revealed spatially heterogenous splicing patterns for MIGs, which was reflected in the proteome, that result in dysregulation of the mutant forelimb’s spatial transcriptome. Using single cell RNAseq, we show that the molecular deficits of the mutant forelimb result in developmental delay of chondroprogenitors. Finally, through 3-dimensional systems reconstructions, we reveal that molecular and cellular stalling of the limb mesenchyme manifests in developmental delay of chondrogenesis and, consequently, disrupted migration of non-endogenous myoblasts and axons. Taken together, our data suggests that upon sensing depletion of LPCs, the limb pauses its molecular program and halts the LPC developmental trajectory, likely affording itself the opportunity to recalibrate prior to furthering development.

## Results

### The U11-null forelimb displays transcriptomic stalling that affects key limb patterning genes

The mutant forelimb experiences a significant reduction in its size that precipitates as early as embryonic day (E)11.5^11^. By P0, the mutant forelimb is drastically reduced in length but maintains proximo-distal segmentation, as it contains a stylopod, a single zeugopod bone, and a single autopodial digit^11^. This phenotype agrees with the symptomology of MOPD1, RS, and LWS, where individuals show reduced limb size but conserved limb patterning^12–15^. Thus, we sought to understand how minor spliceosome inhibition alters the transcriptional trajectory of the developing limb. For this, we interrogated the expression of all protein-coding genes by leveraging our RNAseq datasets of the WT and mutant forelimb at E10.5 and E11.5^11^. We considered a gene expressed if it had transcripts per million (TPM) above 0.5, a TPM slightly below the lowest expression of *Shh* (a lowly-expressed known truth) found among the two WT data sets^18^. With this threshold, we performed pairwise comparisons for each age-matched sample. We found that 57 genes were upregulated (≥1 log2 fold-change (log2FC); *P*≤0.01) and 235 genes were downregulated (≤1 log2FC; *P*≤0.01) in the E10.5 mutant forelimb compared to its WT counterpart (Fig. 1A). At E11.5, 988 genes were upregulated and 15 genes were downregulated in the mutant forelimb (Fig. 1A). Thus, while U11 loss primarily impacts the splicing of MIGs^11^, we secondarily observed a massive transcriptional upregulation by E11.5 (Fig. 1A).

**Figure 1.**
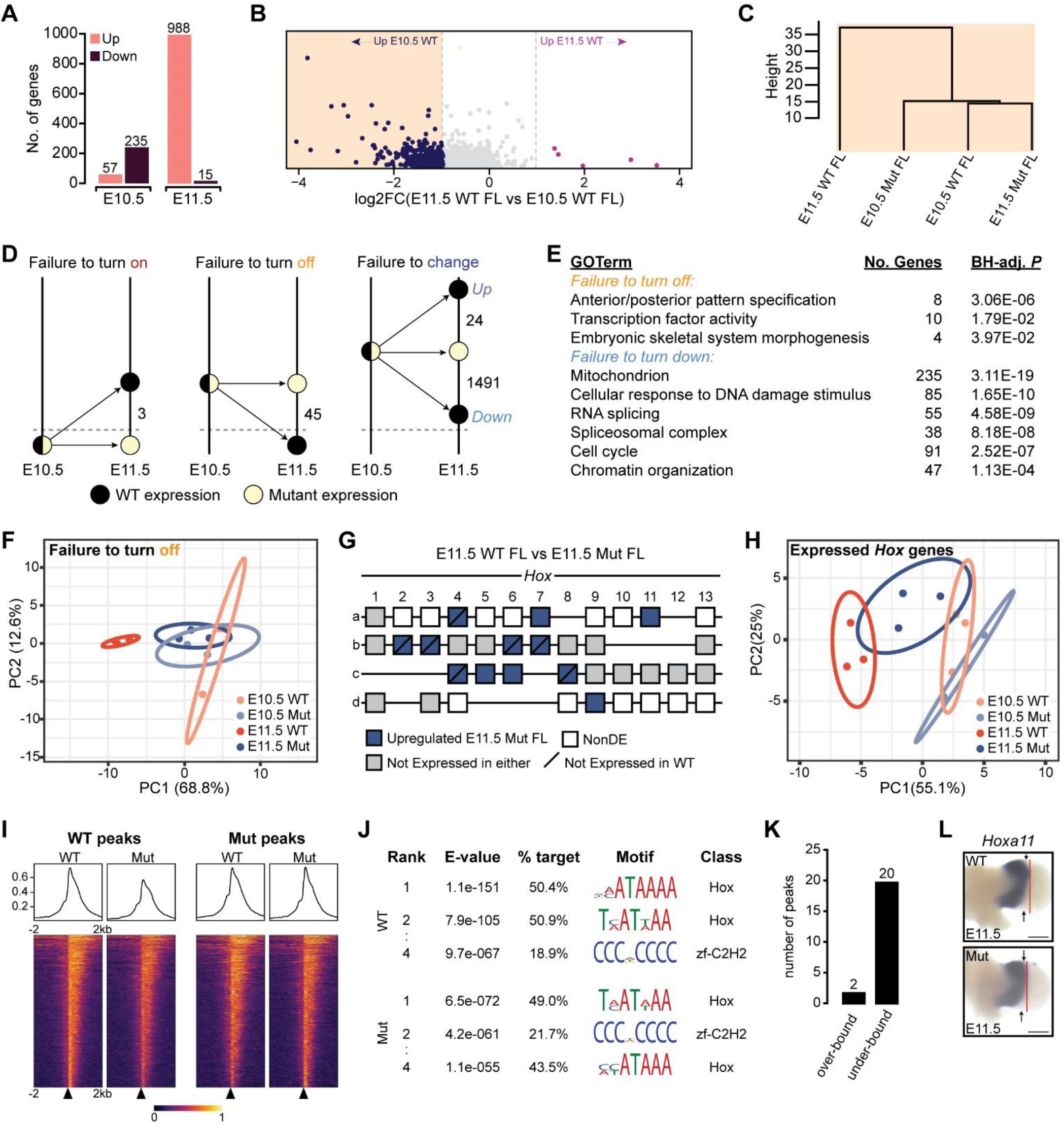
U11 loss results in transcriptomic stalling that affects Hox gene expression. (A) Bar-chart showing number of upregulated and downregulated protein-coding genes in the embryonic day (E) 10.5 and E11.5 mutant forelimb when compared to its appropriate WT control. (B) Scatterplot showing the differential expression of the 988 upregulated protein-coding genes in the E11.5 mutant forelimb from (A) when compared in the WT forelimb (FL) from E10.5 to E11.5. (C) Dendrogram showing the relationship of the subset of upregulated genes in the E11.5 mutant (Mut) forelimb from (A) that are upregulated in the WT forelimb at E10.5 when compared to the E11.5 WT forelimb (dark blue dots in B). (D) Schematics showing the expression patterns of genes stalled in the mutant forelimb. Dashed line shows cutoff for expression. (E) DAVID enrichment for genes that fail to be turned off and genes that fail to be turned down in the mutant forelimb. BH-adj.=Benjamini-Hochberg adjusted. (F) Principal component analysis (PCA) for genes that fail to be turned off in the mutant forelimb. (G) Schematic showing differential expression of *Hox* genes in the E11.5 mutant forelimb compared to the E11.5 WT forelimb. NonDE=non-differentially expressed. (H) PCA for all expressed *Hox* genes in the E10.5 and E11.5 WT and mutant forelimb. Ellipses in PCAs show 95% confidence intervals. (I) Heat-map showing Hoxa11 chromatin-binding profile from −2 kb to 2 kb of transcription start sites (black triangles) in E11.5 WT and mutant forelimb through CUT&RUN. (J) Top three shared Hoxa11 binding targets and their motifs between the E11.5 WT and mutant forelimb. (K) Number of differentially bound peaks in the mutant forelimb at E11.5. (L) Whole mount *in situ* hybridization for *Hoxa11* in E11.5 WT and mutant forelimb. Red line shows distal most signal. Black arrowheads show regions of expanded Hoxa11 expression in mutant forelimb. Scale bar shows 50 μm. Supplemental information for data shown in this figure can be found in Data File 1.

We next sought to understand the WT temporal expression pattern for the 988 upregulated genes in the mutant forelimb. Of these, 311 genes were expressed more highly in the E10.5 WT forelimb, 633 were NonDE, 6 were expressed more highly in the E11.5 WT forelimb, and 38 were not considered expressed in either sample (Fig. 1B). We chose to focus our attention on the 311 genes that were (i) expressed significantly higher in the E11.5 mutant forelimb when compared to the E11.5 WT (Fig. 1A), and (ii) expressed significantly higher in E10.5 WT forelimb when compared to the E11.5 WT (Fig. 1B). This differential expression pattern suggested to us that these genes had transcriptional signatures in the E11.5 mutant forelimb that were reminiscent of an E10.5 WT-like state. To test this, we performed hierarchical clustering of our four datasets using only the expression of these 311 genes (Fig. 1C). Indeed, we found that the E11.5 mutant forelimb clustered most similarly to the E10.5 WT forelimb, leading us to hypothesize that a subset of the mutant transcriptome was developmentally stalled (Fig. 1C).

To capture genes whose expression was developmentally stalled in the mutant forelimb, we defined four categories: (1) genes that failed to be turned on, (2) genes that failed to be turned off, (3) genes that failed to be turned up, and (4) genes that failed to be turned down (Fig. 1D). We analyzed all expressed genes for their presence in one of these bins and discovered that the two most represented sets of genes showing developmental stalling in the mutant forelimb between E10.5 and E11.5 included 45 genes that failed to be turned off and 1,491 genes that failed to be turned down (Fig. 1D). DAVID analysis of genes that had been failed to be turned off revealed enrichment for limb development pathways, including “anterior/posterior pattern specification” and “embryonic skeletal system morphogenesis” (Fig. 1E). Moreover, genes that failed to be turned down enriched for many general biological terms such as “DNA maintenance”, “splicing”, “cell cycle”, and “chromatin organization” (Fig. 1E). We sought to corroborate our developmental stalling binning approach by performing principal component analysis (PCA) for genes that had failed to be turned off. Indeed, PCA revealed that while there was clear separation of the E10.5 and E11.5 WT forelimb datasets, both E10.5 and E11.5 mutant datasets clustered among each other and among the E10.5 WT dataset (Fig. 1F).

Next, we explored individual genes that failed to be turned off. We found that 6 of the 8 genes that enriched for “anterior/posterior pattern specification” were *Hox* genes (Fig. 1E). Thus, we explored transcriptional changes for all *Hox* genes at E11.5. We found that *Hoxa4, a7, a11, Hoxb2, b3, b4, b5, b6, b7, Hoxc4, c5, c6, c8*, and *Hoxd9* were upregulated, while none were downregulated in the mutant forelimb (Fig. 1G). Through PCA, we found that the E11.5 mutant forelimb possesses a *Hox* transcriptional program that bridges the E10.5 and E11.5 WT state (Fig. 1H). Given that *Hox* genes regulate limb patterning^19^, we proposed their mis-expression would alter their function and potentially contribute to micromelia upon U11 loss^11^. One notable upregulated gene in this list is *Hoxa11* as it is a critical regulator of zeugopod formation^20^, which we have previously reported is disproportionately reduced in the P0 U11-null limb^11^. Thus, we hypothesized that Hoxa11 may have an aberrant DNA binding footprint in the mutant forelimb. To test this, we performed cleavage under targets and release using nuclease (CUT&RUN)^21^ for Hoxa11 at E11.5 (Fig. 1I). Hoxa11 was bound to similar motifs in the mutant forelimb as compared to the WT (Fig. 1J). However, we found that the frequency of Hoxa11 bound to targets with the top motif identified in the WT dataset was reduced by ~13% (Fig. 1J). In addition, we found 22 differentially bound peaks, with two peaks over-bound in the mutant forelimb and 20 peaks under-bound in the mutant forelimb (Fig. 1K). The two over-bound peaks corresponded to promoter regions of *Prrx1*, a marker of early limb progenitor cells^17^, and *Sin3a*, a transcriptional repressor that regulates cell proliferation^22^ (Fig. 1K). Amongst the 20 under-bound peaks was *Noc2l*, which is known to repress p53-mediated apoptosis^23^ (Fig. 1K). Lastly, we confirmed the upregulation of *Hoxa11* in the mutant forelimb through whole mount in situ hybridization (WISH), which revealed that *Hoxa11* expression expanded anteriorly and posteriorly along the distal edge of the mutant forelimb (Fig. 1L). This shift in the *Hoxa11* expression pattern suggested that transcriptomic alterations are spatially biased in the U11-null forelimb.

### Microdomain RNAseq reveals spatially heterogenous mis-splicing of MIGs in the U11-null forelimb

Distal posterior expansion of *Hoxa11* expression reinforced our previous findings that the consequences of U11 loss are spatially biased, such as observed for cell death, which we had found to be distal posteriorly concentrated^11^. To confirm this phenomenon, we performed immunofluorescence for cleaved caspase 3 (CC3), an early marker of apoptosis, on serial sections of an E11.5 WT and mutant forelimb. By 3-dimensionally reconstructing this signal, we found that the highest concentration of CC3-positive cells was in the distal posterior region, followed by the distal anterior region (Fig. 2A). These observations suggested to us that either i) U11 loss causes the same splicing defects throughout the limb, but spatially distinct populations of limb progenitor cells respond differently to these deficits; ii) U11 loss leads to distinct splicing defects in different domains of the limb; or iii) both. To distinguish between these hypotheses, we harvested N=3 WT and mutant embryos at E11.5 and micro-dissected the forelimb into four regions – proximal anterior (PA), proximal posterior (PP), distal anterior (DA), and distal posterior (DP) – and subsequently performed total RNAseq (Fig. 2B). We first confirmed downregulation of *Rnu11* in the mutant microdomains, which we found was significant for DA, DP, and PA, but not PP (Fig. 2C).

**Figure 2.**
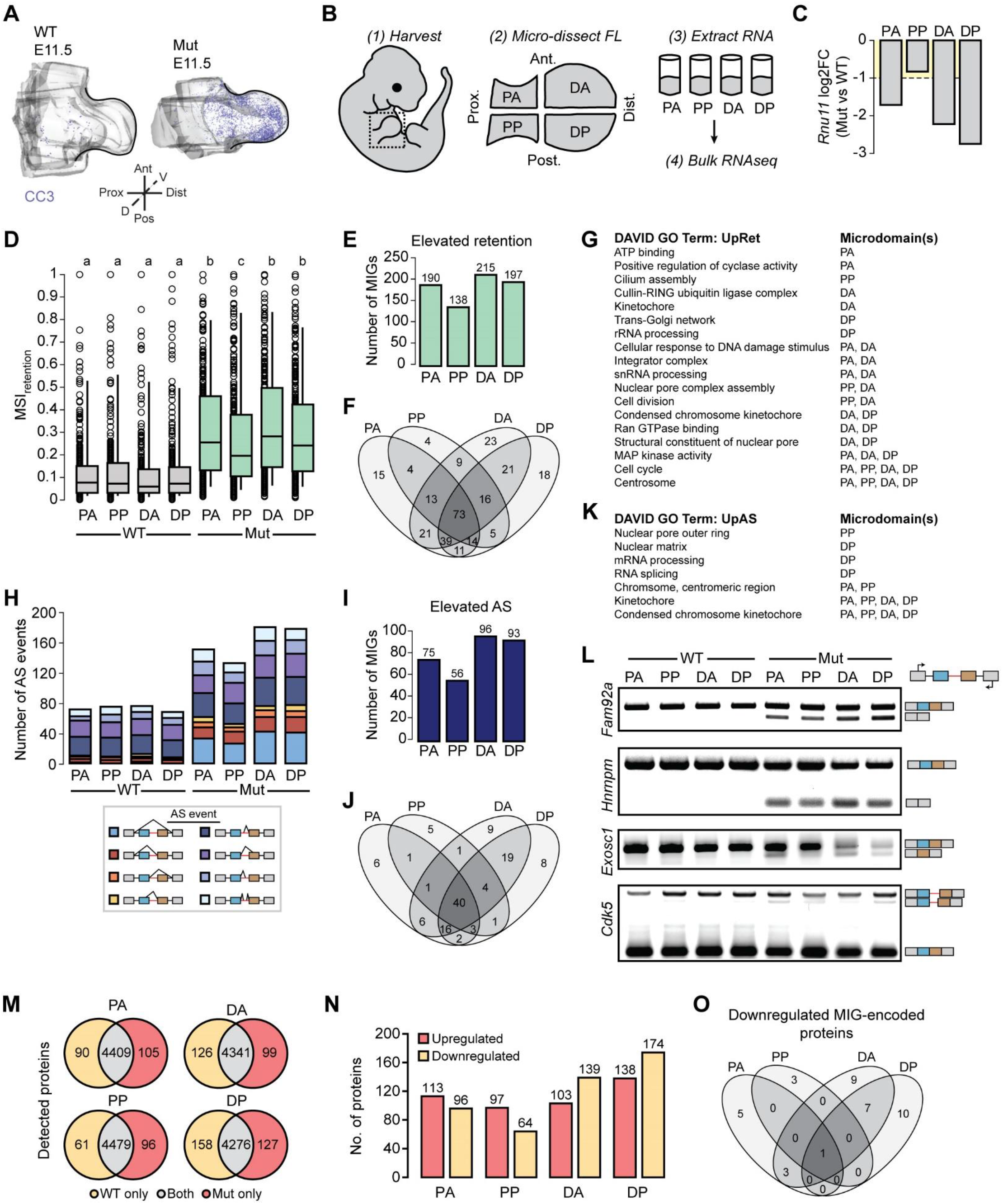
U11 loss results in spatially heterogenous mis-splicing of minor introns. (A) Three-dimensional reconstruction of CC3+ cells in the E11.5 WT and mutant (Mut) forelimb determined through serial immunofluorescence. Prox=proximal, Dist=distal, Ant=anterior, Pos=posterior, D=dorsal, V=ventral. (B) Schematic showing micro-dissection workflow. FL=forelimb; PA=proximal anterior; PP=proximal posterior; DA=distal anterior; DP=distal posterior. (C) Differential expression (log2 fold-change) of *Rnu11* in mutant (mut) microdomains when compared to corresponding WT counterpart as determined by RNAseq. Dashed line shows cut-off for significance. (D) Box plot showing 10th to 90th percentile mis-splicing index for 385 minor introns showing varying levels of minor intron retention in all eight samples. Samples with different letters have significantly different median MSIs (*P*<0.05) as determined by all-to-all Kruskal-Wallis test with *post-hoc* Dunn’s test. (E) Bar chart showing number of MIGs with significantly elevated minor intron retention in each mutant microdomain when compared to its appropriate WT counterpart. (F) Overlap analysis for MIGs with elevated minor intron retention in each mutant microdomain. (G) Gene ontology (GO) terms identified for MIGs with significantly upregulated minor intron retention (UpRet) in each individual mutant microdomain (left) with each microdomain showing enrichment for a given term listed (right). (H) Bar chart showing number of detected alternative splicing (AS) events in each mutant microdomain. (I) Bar chart showing number of AS events significantly upregulated in each mutant microdomain when compared to its appropriate WT counterpart. (J) Overlap analysis for MIGs with upregulated AS in each mutant microdomain. (K) GO terms enriched by MIGs with significantly upregulated AS (UpAS) in each individual mutant microdomain (left) with each microdomain showing enrichment for a given term listed (right). (L) RT-PCR validations of AS events detected to be significantly increased in mutant microdomains for *Fam92a, Hnrnpm, Exosc1*, and *Cdk5*. Red line shows minor intron. Black lines show major introns. Light blue box shows 5’ upstream exon. Brown box shows 3’ downstream exon. Black arrows show primer location for each interrogated MIG. Predicted spliced products shown adjacent to their corresponding gel band. (M) Venn-diagrams (not to scale) showing the number of detected proteins in each WT and mutant microdomain. (N) Bar-chart showing the number of differentially expressed proteins in each mutant microdomain when compared to its appropriate WT counterpart. (O) Overlap analysis for downregulated MIG-encoded proteins in each mutant microdomain. Supplemental information for data shown in this figure can be found in Data File 2.

To test the possibility that there was differential minor spliceosome inhibition in the mutant microdomains (as read out by changes in minor intron splicing), we quantified a mis-splicing index (MSI) for each microdomain, which measures the amount of minor intron retention relative to canonical splicing for each minor intron. Indeed, each mutant microdomain was found to have a significant increase in minor intron retention relative to its WT counterpart (Fig. 2D). In line with the differential loss of U11, we found that the mutant DA, DP, and PA microdomains showed higher levels of minor intron retention than the mutant PP microdomain (Fig. 2D).

We next identified the MIGs whose minor introns showed significantly elevated retention in each mutant microdomain when compared to their WT counterpart. This yielded 190, 138, 215, and 197 MIGs with significantly elevated minor intron retention in the mutant PA, PP, DA, and DP microdomains, respectively (Fig. 2E). Whereas 73 MIGs showed significantly elevated minor intron retention in all mutant microdomains, we identified an abundance of MIGs with elevated minor intron retention in all potential sample combinations (Fig. 2F). We subsequently leveraged DAVID to identify the biological pathways enriched by MIGs with elevated minor intron retention in each mutant microdomain. In line with the global cell cycle defects we previously reported in the U11-null forelimb^11^, we found that the gene ontology (GO)-terms “cell cycle” and “centrosome” were enriched by MIGs with elevated minor intron retention in all mutant microdomains (Fig. 2G) We also found that each mutant microdomain, as well as combinations of mutant microdomains, enriched for unique GO-terms, such as “Cullin-RING ubiquitin ligase complex” only in mutant DA or “cilium assembly” only in mutant PP (Fig. 2G).

We have previously reported that inhibition of the minor spliceosome results in alternative splicing (AS) around minor introns^24,25^. Therefore, we employed our previously reported minor intron AS bioinformatics pipelines, which employ BEDTools to bin AS events into one of eight categories (including exon skipping, cryptic 5’ splice site usage, cryptic 3’ splice site usage, and cryptic exon usage), to detect the number of AS events occurring in each mutant microdomain^24^. In line with our hypothesis, we found an increase in the number of AS events in each mutant microdomain, which were concentrated mainly in exon skipping and cryptic 5’ splice site events (Fig. 2H). Altogether, we quantified a significant increase in 75, 56, 96, and 93 AS events in the mutant PA, PP, DA, and PP microdomains, respectively, when compared to their appropriate WT counterpart (Fig. 2I). Whereas 40 MIGs showed significant AS in all four mutant microdomains, we nonetheless detected unique MIGs with AS in each possible sample combination (Fig. 2J). Consistent with our minor intron retention findings, MIGs with elevated AS in all mutant microdomains enriched for “condensed chromosome kinetochore”, further supporting the global cell cycle defect experienced throughout the U11-null forelimb^11^ (Fig. 2K). Moreover, overlap analysis of DAVID GO-terms generated by MIGs with significantly elevated AS for each mutant microdomain revealed enrichment for unique terms from mis-spliced MIGs in the PP, DP, and PA+PP (Fig. 2K).

Through RT-PCR, we validated multiple shared and unique AS events among the four mutant microdomains. We found robust exon skipping for both exons flanking the minor intron in *Fam92a* (also known as *Cibar1*), which encodes a critical regulator of hedgehog signaling in the developing limb^26,27^ (Fig. 2L). We validated the same category of AS detected for *Hnrnpm*, which encodes an essential, ubiquitously expressed splicing factor whose genetic ablation in mice leads to pre-weaning lethality^28–30^ (Fig. 2L). Notably, our RNAseq analysis quantified this AS event to be significantly elevated in the distal mutant microdomains when compared to the proximal mutant microdomains, which was captured by RT-PCR (Fig. 2L). We next validated an upstream exon skipping event for *Exosc1* that our RNAseq analysis detected in all mutant microdomains except PP, and a cryptic 3’ splice site usage for *Cdk5* that our RNAseq analysis detected in all mutant microdomains except DA (Fig. 2L). *Exosc1* is a crucial member of the exosome RNA surveillance complex and its loss in mice also leads to pre-weaning lethality^30–32^. Moreover, *Cdk5* encodes a kinase that regulates nervous system development through various pathways, including neuronal differentiation, axonal migration, and synaptogenesis^33^.

Mis-splicing of MIGs is predicted to result in downregulated expression of their endogenously encoded proteins. Thus, we performed LC-MS/MS on microdomains collected from E11.5 WT and mutant forelimbs. We detected >4,400 proteins within each microdomain, of which >96% were shared between both conditions per microdomain (Fig. 2M). Nonetheless, each microdomain contained sets of proteins detected only in the WT or only in mutant, which we deemed as downregulated or upregulated, respectively (Fig. 2M, N). In addition, for proteins that were detected in both WT and mutant samples per microdomain, we employed student’s two-tailed T-test with Benjamini-Hochberg *P*-value adjustment to identify those expressed significantly higher or lower in the mutant microdomain (BH < 0.05) (Supp. Fig. 1). We did not incorporate a fold-change cut-off to avoid omitting potential false-negatives. Ultimately, we grouped the uniquely detected proteins with those significantly differentially expressed into their respected upregulated or downregulated bins for each microdomain (Fig. 2N). With this approach, we discovered 113, 97, 103, and 136 proteins upregulated in the mutant PA, PP, DA, and DP, respectively (Fig. 2N). Additionally, we found 96, 64, 139, and 174 proteins downregulated in the mutant PA, PP, DA, and DP, respectively (Fig. 2N). Overlap analysis of the downregulated MIG-encoded proteins revealed a high degree of heterogeneity, as only a single MIG-encoded protein (Pdpk1) was downregulated in all four mutant microdomains (Fig. 2O). Notably, 5 MIG-encoded proteins were downregulated only in the mutant PA microdomain, 3 in PP, 9 in DA, and 10 in DP, whereas 3 were shared to be downregulated anteriorly (PA+DA) and 7 were shared to be downregulated distally (DA+DP) (Fig. 2O). This data corroborated that spatial molecular heterogeneity regarding mis-splicing of MIGs upon U11 loss manifests in regionally biased downregulation for MIG-encoded proteins.

### Spatially distinct splicing patterns can be linked to spatial gene expression changes in the U11-null forelimb

Given the abundance of regionally specific mis-splicing events identified through our microdomain approach, we hypothesized that U11 loss would drive spatially distinct changes in gene expression. To test this hypothesis, we performed a head-to-head differential gene expression analysis for each WT and mutant microdomain pair (ex: WT-DA vs Mut-DA) through isoEM2 and isoDE2^34^. Using a 1 TPM threshold for gene expression, we found an abundance of upregulated genes in each microdomain with few downregulated genes (Supp. Fig. 2A-B). Overlap analysis for the number of differentially expressed genes for each possible combination of microdomains yielded a high degree of dissimilarity, with the greatest overlap found for upregulated genes (53) shared by both distal mutant microdomains (Supp. Fig. 2A-B). Indeed, we identified only nine genes (including one MIG: *Mapk8*) upregulated in all four mutant microdomains and only two genes (including one MIG: *Phyh*) downregulated in all four mutant microdomains (Supp. Fig. 2A-B). In line with this heterogeneity, we found that the biological pathways enriched by the differentially expressed genes for each microdomain were mostly distinct (Supp. Fig. 2C-D).

Since we identified regional heterogeneity for changes in both minor intron splicing and differential gene expression, we hypothesized that microdomain-specific gene expression changes were influenced by microdomain-specific splicing alterations. One way this could happen is if these genes were part of shared biological networks. To test this idea, we employed STRING to identify known interactions between genes underlying regionally enriched biological pathways and MIGs with microdomain-specific upregulation of minor intron retention or AS. For example, we found that submission of the eight genes that were upregulated in the mutant DA microdomain and enriched for KEGG “p53 signaling pathway” (Supp. Fig. 2C) along with the 23 and 9 MIGs showing uniquely upregulated minor intron retention and AS, respectively, in this microdomain (Fig. 2F, J) generated a tight-knit network, connecting all upregulated genes with a subset of the mis-spliced MIGs (Supp. Fig. 2E). Notably, we found that the 18 and 8 MIGs with uniquely upregulated minor intron retention and AS, respectively, in the mutant DP microdomain (Fig. 2F, J) showed distinct connectivity patterns when submitted to STRING with the DP-specific upregulated genes underlying the GO terms “extracellular matrix structural constituent” (Supp. Fig. 2D, F) or “ATP binding” (Supp. Fig. 2D, G). Whereas 10 mis-spliced MIGs formed a concise network with nine of the “extracellular matrix structural constituent” genes, they did so by all feeding into one single MIG – *Eed* – that connected at multiple nodes in the upregulated gene network (Supp. Fig. 2F). On the other hand, we found that 16 mis-spliced MIGs were integrated throughout a network with 32 “ATP-binding genes” (Supp. Fig. 2G). In either case, disruption in the expression or function of the encoded MIG product upon mis-splicing is predicted to dramatically affect the functions of these networks. Thus, our findings suggest that regionally biased mis-splicing causes disrupted gene expression regulation for non-MIGs, resulting in spatially distinct gene expression changes.

### Spatial transcriptomic patterns governing the transition from limb patterning to differentiation are stalled upon U11 loss

The developing limb relies upon spatially discrete gene expression profiles for its patterning^35^. Therefore, detection of spatially distinct changes in splicing and gene expression in the mutant forelimb led us to suspect that this would lead to alterations in its spatial transcriptome. To test this, we needed to devise a method to generate a spatial atlas of gene expression for the WT forelimb, from which we could compare whether the mutant successfully established the appropriate spatial gene expression profiles. Ultimately, we leveraged our microdomain RNAseq datasets and approximated each gene’s expression pattern in the developing limb by generating what we refer to as a *spatialPlot*, a graph that has predictive value for the location of a gene’s expression within a tissue. Using a 1 TPM cut-off for expression, we omitted all genes found to be below this threshold in all four microdomains. Next, we averaged each expressed gene’s TPM values among proximal domains (PA, PP), distal domains (DA, DP), anterior domains (PA, DA), and posterior domains (PP, DP). With these averaged values, we calculated each gene’s proximo-distal log2FC and antero-posterior log2FC and plotted them such that their graphical position corresponds to their anticipated spatial position in the developing E11.5 WT forelimb (Fig. 3A). We then highlighted genes that were statistically significantly enriched in their respective domain as determined by ANOVA followed by post-hoc Tukey’s test (Fig. 3A). For example, *Pax1* was found to have a proximo-distal log2FC of −7.3 and an antero-posterior log2FC of 4.4, positioning it proximal-anteriorly on the *spatialPlot* (Fig. 3A). Indeed, it has been shown through whole mount *in situ* hybridization (WISH) that *Pax1* expression is restricted to the PA domain in the E11.5 WT forelimb^36^. Importantly, we found spatially enriched expression for previously validated genes through WISH for each region, including *Pkdcc* proximally^37^, *Dlk1* in PP^38^, *Hand2* posteriorly^39^, *Shh* in DP^18^, *Hoxa13* distally^40^, *Zic3* in DA^41^, and *Asb4* anteriorly^42^ (Fig. 3A). We further confirmed the validity of this approach by identifying GO terms enriched by each of the spatial gene sets (Supp. Fig. 3A). Distally restricted genes enriched for terms such as “proximo-distal pattern formation” and “positive regulation of cell proliferation”, whereas proximally restricted genes enriched for “skeletal system development”, “nervous system development”, “skeletal muscle tissue development”, and “regulation of cell migration” (Supp. Fig. 3A). Notably, PA genes enriched for terms including “chondrocyte differentiation”, whereas PP genes only enriched for “proteinaceous extracellular matrix”, which is indicative of chondrogenesis occurring in the PA region but not yet as robustly in the PP region at E11.5^43^ (Supp. Fig. 3A). In addition, we found enrichment for “hedgehog receptor activity” and other pathways downstream of Shh signaling in the DP enriched genes (Supp. Fig. 3A).

**Figure 3.**
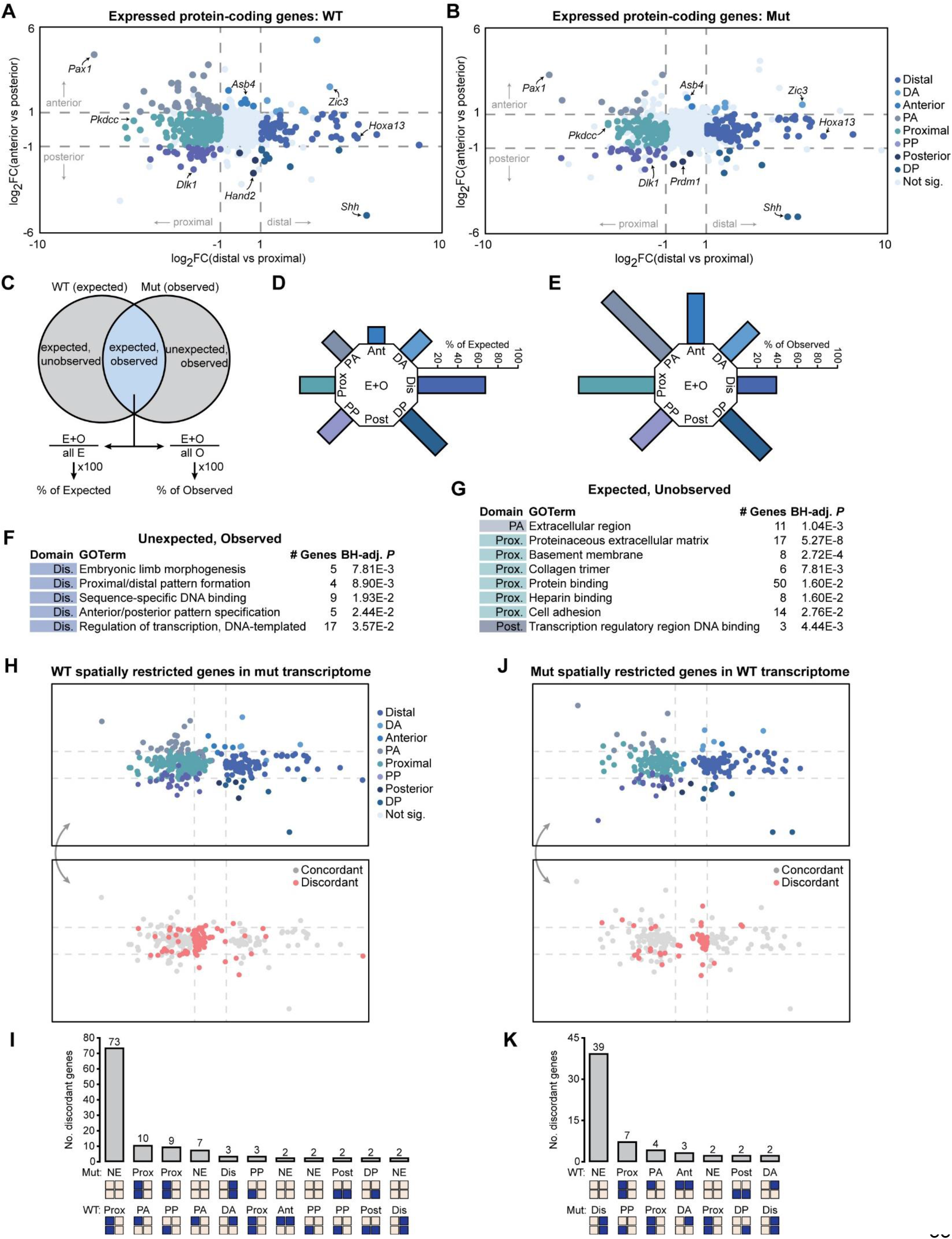
U11 loss results in failure to separate proximal differentiation genes from distal patterning genes. (A-B) *spatialPlot* for expressed protein-coding genes in E11.5 WT (A) and mutant (B) forelimb. Dashed lines show cut-off for spatial enrichment. Labeled genes show known-truths. (C) Schematic showing mathematical workflow for determining percentage of expected yet unobserved genes and observed yet unexpected genes in the mutant *spatialPlot*. (D) Wrapped-around bar chart showing percentage of mutant spatially enriched genes that were expected and observed out of all expected genes (based on their spatial enrichment in WT) for each region. (E) Wrapped-around bar-chart showing percentage of mutant spatially enriched genes that were expected and observed out of all observed genes (based on their spatial enrichment in mutant) for each region. (F-G) Gene ontology (GO) enrichment for unexpected yet observed genes (F) and expected yet unobserved genes (G) with corresponding domain in which those genes were found to be spatially enriched. Prox=proximal; dis=distal; ant=anterior; post=posterior; PA=proximal anterior; PP=proximal posterior; DA=distal anterior; DP=distal posterior. (H) *spatialPlot* generated by graphing WT spatially enriched genes using their expression values in the mutant forelimb (top), with concordance, i.e., correct matching of spatial enrichment in both WT and mutant transcriptome, superimposed (bottom). (I) Bar chart depicting patterns of expression discordance, showing only those with more than one gene. Simplified limb bud schematics shown below depict respective category of regional enrichment. Blue boxes show expression enrichment, light yellow boxes show lack of expression enrichment. NE=not enriched in any region. (J) *spatialPlot* generated by graphing Mut spatially enriched genes using their expression values in the WT limb (top), with concordance superimposed (bottom). (K) Bar chart depicting patterns of expression discordance, showing only those with more than one gene. Prox=proximal; Dis=distal; Ant=anterior; Post=posterior; PA=proximal anterior; PP=proximal posterior; DA=distal anterior; DP=distal posterior. Supplemental information for data shown in this figure can be found in Data File 3.

We then employed the same analysis on the E11.5 mutant forelimb to test the hypothesis that its spatial transcriptome was perturbed. We previously showed that the expression of select genes of the core limb patterning network was spatially maintained in the E11.5 U11-null limb. Thus, we were first able to validate our dissection integrity by benchmarking the RNAseq data with expression of known-truths used for the WT *spatialPlot*, aside from *Hand2*, which was enriched in the mutant DP domain rather than posterior domain (Fig. 3B). Nonetheless, we found *Prdm1* was enriched in the mutant posterior domain, which has been previously shown to be expressed in the posterior WT forelimb at E11.5 by WISH^44^ (note: *Prdm1* was enriched in WT posterior by FC, but not by ANOVA with BH-adjustment (*P*=0.06), likely reflecting a false negative due to our stringent thresholding) (Fig. 3B). Moreover, we found similar biological pathway enrichments as observed for the WT upon submitting spatially restricted genes in the mutant forelimb to DAVID (Supp. Fig. 3B). Thus, the mutant *spatialPlot* confirmed that the core limb patterning network had proceeded sufficiently by E11.5 to establish fundamental gene expression separation, such as positioning *Shh* in the DP domain (Fig. 3B).

We next sought to determine how well the total mutant spatial transcriptome reflected that of the WT. For this, we treated the genes found to be enriched in each WT microdomain as “expected” genes, and genes found to be enriched in each mutant microdomain as “observed” genes (Fig. 3C). We then identified the number of enriched genes in each microdomain that were found in the WT only (expected yet unobserved), in the mutant only (unexpected yet observed), and in both (expected and observed) (Fig. 3C). Subsequently, we used these numbers to calculate the percentage of genes that were expected and observed out of the total expected genes for each microdomain (Fig. 3C). In addition, we calculated the percentage of genes that were expected and observed out of the total observed genes for each microdomain (Fig. 3C). The former analysis allowed us to determine the proportion of genes within a mutant microdomain that matched the WT spatial transcriptome, whereas the latter allowed us to determine the abundance of genes within a mutant microdomain that were uniquely enriched in the mutant. For example, whereas the distally enriched genes in the mutant represented 67% of expected genes, we found that this was only 39% of the total observed distally enriched genes (Fig. 3D-E). Thus, there was an abundance of unexpected yet observed genes enriched distally in the mutant (Fig. 3E). In contrast, we found that while proximally enriched genes in the mutant forelimb represented only 35% of expected proximally restricted genes, this number nonetheless corresponded to 75.5% of all observed proximally restricted genes (Fig. 3D-E). Thus, there was an abundance of expected yet unobserved genes proximally in the mutant. Of note is that three genes were identified to be enriched posteriorly in both the WT and mutant forelimb, yet none of these were shared (Fig. 3D-E).

We next used DAVID to identify biological pathways that were present among genes either unexpected yet observed or expected and unobserved for each mutant microdomain. Of genes unexpected yet observed, only those found distally had biological enrichment, yielding GO terms including “embryonic limb morphogenesis”, “proximal/distal pattern formation”, and “anterior/posterior pattern formation” emerged (Fig. 3F). On the other hand, expected and unobserved genes among the mutant PA, proximal, and posterior microdomains enriched for many terms of differentiation, including “proteinaceous extracellular matrix” and “collagen trimer” (Fig. 3G). This suggested to us that the spatial transcriptome of the E11.5 mutant forelimb failed to appropriately separate its early limb patterning program from its subsequent differentiation program.

Next, we aimed to decipher the exact spatial changes underlying the mutant forelimb’s disrupted spatial transcriptome. Thus, we extracted all spatially restricted genes found in the WT forelimb and identified their expression patterns in the mutant forelimb. Subsequently, we generated a new *spatialPlot*, from which we determined whether each WT spatially enriched gene showed the same spatial expression pattern in the mutant forelimb (Fig. 3H). If it did, we deemed this gene concordant; however, if it was enriched in a different region, or not enriched in any region, it was deemed discordant (Fig. 3H). Our analysis identified a dramatic spatial bias in discordant gene expression, with 73 proximally restricted genes in the WT showing no spatial enrichment in the mutant transcriptome (Fig. 3I). We then aimed to determine whether the 5 MIGs with uniquely upregulated minor intron retention or AS events occurring in the mutant proximal microdomains might underlie this discordance (PA+PP in Fig. 2F, J). Thus, we submitted these 78 genes to STRING and identified the largest network generated, which connected 24 proximally discordant genes with one MIG, *Raf1*, that encodes the Raf-1 serine/threonine kinase (Supp. Fig. 4A). The importance of *Raf1* in this network is underscored by the fact that mutations in it have been linked to Noonan Syndrome, which is characterized by short stature^45^. This phenotype is not surprising considering that this network contains multiple genes essential for skeletogenesis, including *Shox2*^46–50^ and *Col2a1*^51–53^ (Supp. Fig. 4A). Indeed, loss of *Col2a1* in mice results in severe skeletal dysplasia^51^. Moreover, *Shox2*-null mice show dramatic reduction in stylopod length, as well as deficits in the migration of axonal projections and muscle precursors into the proximal forelimb, suggesting that it integrates the development of multiple systems during limb organogenesis^46–50^. Altogether, disrupted splicing of transcripts emanating from *Raf1* could perturb the appropriate development of the skeletal, nervous, and muscular systems.

Equally important to genes dis-enriched in the mutant transcriptome are those that are uniquely enriched. Thus, we performed the same analysis as described above in reverse, wherein we extracted all spatially restricted genes found in the mutant forelimb, identified their expression patterns in the WT transcriptome, and generated a novel spatial plot to determine concordance and discordance (Fig. 3J). Again, we found a substantial spatial bias in discordance such that 39 genes distally enriched in the mutant transcriptome were not spatially enriched in the WT transcriptome (Fig. 3K). In turn, we submitted these genes, along with the 40 MIGs with uniquely upregulated minor intron retention or AS events found in the mutant distal microdomains (DA+DP in Fig. 2F, J) to STRING. This approach generated a tight-knit network containing 10 mis-spliced MIGs interacting with 11 distally discordant genes (Supp. Fig. 4B). Interestingly, this network contains two genes, *Etv4* and *Dusp6*, essential for limb development due to their regulation of Fgf signaling and, in turn, Shh activity, indicating that distally mis-spliced MIGs are implicated in regulating the transcriptional integrity of distal limb patterning networks^54,55^ (Supp. Fig. 4B).

### Single cell RNAseq reveals developmental delay of chondroprogenitors in the U11-null forelimb

One possibility underlying the molecular failure of the U11-null forelimb to separate its proximal differentiation program from its distal patterning program is that it contains altered abundances of progenitor cell populations governing these processes. To test this, we performed single cell RNAseq on pooled (N=6) E11.5 WT and mutant forelimbs. We captured 24,516 high-quality WT cells (median genes/cell: 2,621; median transcripts/cell: 8,056.5) and 31,038 high-quality mutant cells (median genes/cell: 1,934, median transcripts/cell: 5,250) (Supp. Fig. 5). UMAP-mediated clustering of the WT cells yielded 16 distinct populations: nine clusters of endogenous limb mesenchymal cells (ELM; *Prrx1+, Sox9+;* clusters 0-8; 21,497 cells), two clusters of muscle progenitors (*Myod1+;* clusters 9-10; 1,794 cells), one skin cluster (*Krt14+;* cluster 12; 325 cells), one vasculature cluster (*Cdh5+;* cluster 13; 209 cells), two blood clusters (*Hbb-a1+, Lst1+;* clusters 11, 14; 626 cells), and one cluster of neural endothelial cells (*Phactr1+;* cluster 15; 65 cells)^56^ (Fig. 4A-B). Separate yet identical analysis of the mutant cells yielded 15 clusters: eight clusters of ELM (clusters 0-3, 5-8; 25,134 cells), two clusters of muscle progenitors (clusters 4, 9; 3,802 cells), one cluster of skin progenitors (cluster 11; 535 cells), one cluster of vasculature cells (cluster 12; 442 cells), two clusters of blood cells (clusters 10, 13; 1,039 cells), and one cluster of neural endothelial cells (cluster 14; 86 cells) (Fig. 4C-D). We next sought to determine whether the abundances of these broad cell classes were maintained in the U11-null forelimb. Whereas we found that 87.7% of the WT cells were classified as ELM, only 81.0% of mutant cells were ELM (FC=0.92) (Fig. 4E-F). In turn, the mutant forelimb showed increased abundances of all non-ELM cell types, with the greatest increase being observed for vasculature (0.9% vs 1.4%; FC=1.67) and muscle progenitors (7.3% vs 12.3%; FC=1.67) (Fig. 4E-F). This decrease in mutant ELM is in line with significantly elevated apoptosis at this time point (Fig. 2A).

**Figure 4.**
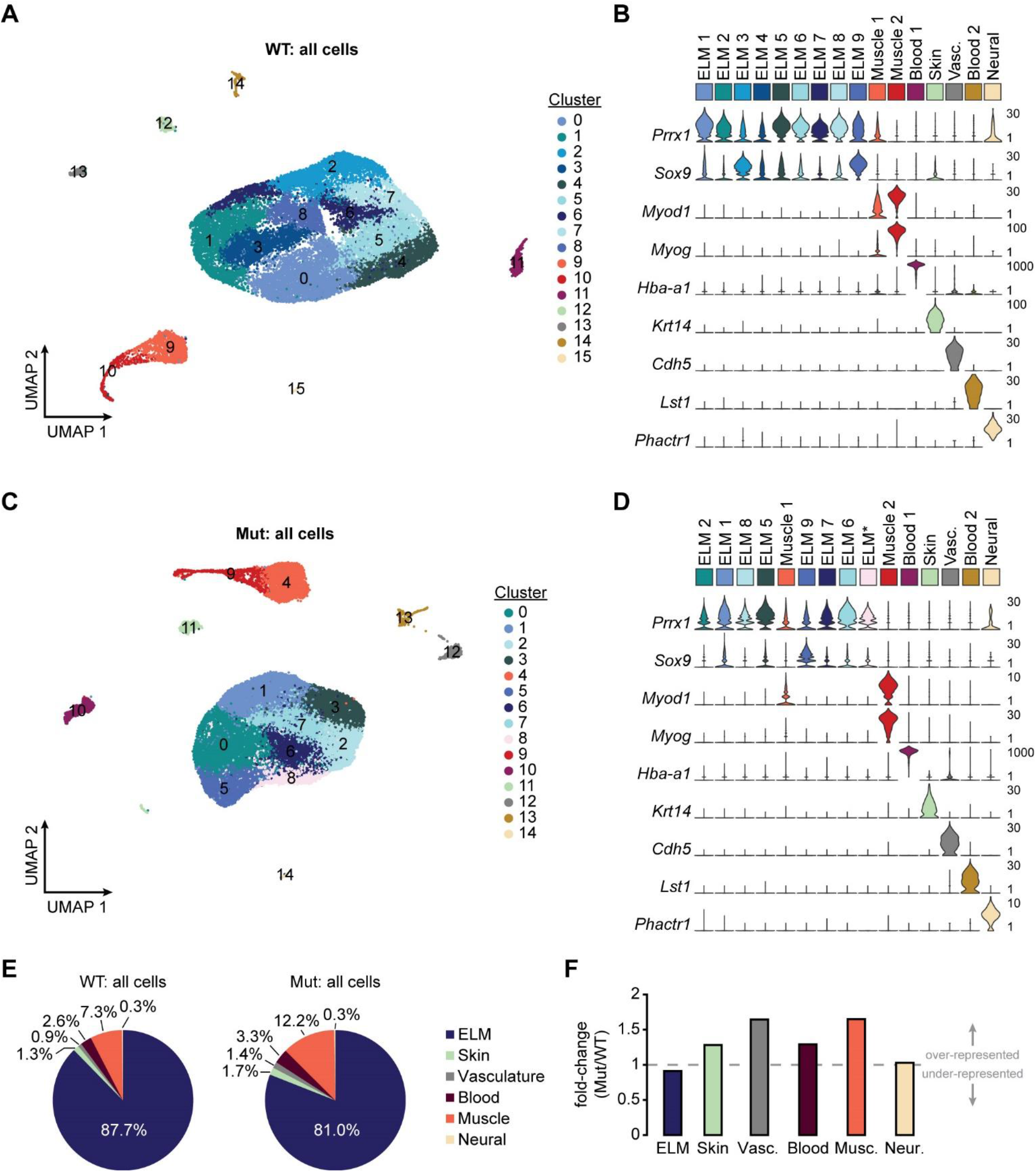
Altered cell type abundances in the U11-null forelimb. (A-D) UMAP showing the sixteen identified WT (A) and fifteen identified mutant (C) cell clusters at E11.5, with violin plots showing gene expression profiles used for WT (B) and mutant (D) cluster classification. Mutant clusters (C-D) were matched back to their closest WT counterpart (A-B) using the top 20 identified transcriptional markers for each cell population. (E) Pie chart showing the distribution of all identified cells in the WT (left) and mutant (right) forelimb. Similar cell types were grouped into broader classes. (F) Bar chart showing the fold-change for the abundance of each broad cell class in the mutant forelimb relative to the WT forelimb. Values were calculated by dividing the percentage abundance in the mutant by the percentage abundance in the WT. Dashed line shows fold-change=1. ELM=endogenous limb mesenchyme; Vasc.=vasculature; Neur.=neural endothelial cells, *=identified only in mutant.

Whereas we found equal numbers of non-ELM clusters with similar transcriptional markers, we found that the mutant forelimb lacked two ELM clusters identified in the WT and contained one unique ELM cluster (Fig. 4B, D). Thus, we sought to integrate the two ELM datasets such that we could interrogate their cellular distribution simultaneously and, in turn, directly determine changes in cell-type abundances. UMAP-mediated clustering of integrated WT and mutant ELM yielded 13 clusters, of which cells from both the WT and mutant forelimb were present in each cluster (Fig. 5A). As our current and previous data show that the molecular and cellular response to U11 loss is spatially heterogenous^11^, we hypothesized that changes in ELM cell type abundances would be spatially heterogenous. We therefore aimed to determine the relative position and differentiation state of each WT and mutant ELM cell. To determine cell position, we leveraged our microdomain RNAseq data, wherein we were able to define signature genes for each microdomain (Supp. Fig. 6). By using the top eight identified signature genes for each microdomain as modules with Seurat’s addModuleScore, we could determine the similarity of every cell to the four microdomains (Fig. 5B). Utilizing a strategy similar to our *spatialPlot*, we binned every cell into one of nine spatial locations (DA, DP, PA, PP, proximal, distal, anterior, posterior, or center) based on the difference between their scores along the proximo-distal axis (distal score – proximal score) and antero-posterior axis (anterior score – posterior score) (Fig. 5C). We then projected this into a *spatialPlot* and superimposed cluster identity for both the WT and mutant forelimb (Fig. 5D-E). The integrity of our binning strategy was confirmed through expression of known regional markers in their corresponding domain. For example, *Asb4* expression was specific to cells binned in PA, anterior, and DA microdomains^42^ (Supp. Fig. 7). Similarly, *Pax1*, a known marker of the PA microdomain^36^, was uniquely expressed by cells only binned in the PA microdomain (Supp. Fig. 7). Finally, *Pmaip1*, a gene with novel spatial expression found through our microdomain analysis, was exclusively expressed by cells in the DA and distal microdomains (Supp. Fig. 7). After determining each cell’s approximate spatial location, we approximated each cell’s differentiation state using previously reported gene expression markers (limb progenitor cell (LPC): *Pdgfra+, Sox9-, Col2a1-*; osteochondroprogenitor (OCP): *Sox9*+, *Col2a1-*; early-stage chondroblast (CB) *Col2a1+*, low; late-stage CB: *Col2a1+*, high)^57^ (Fig. 5F-G).

**Figure 5.**
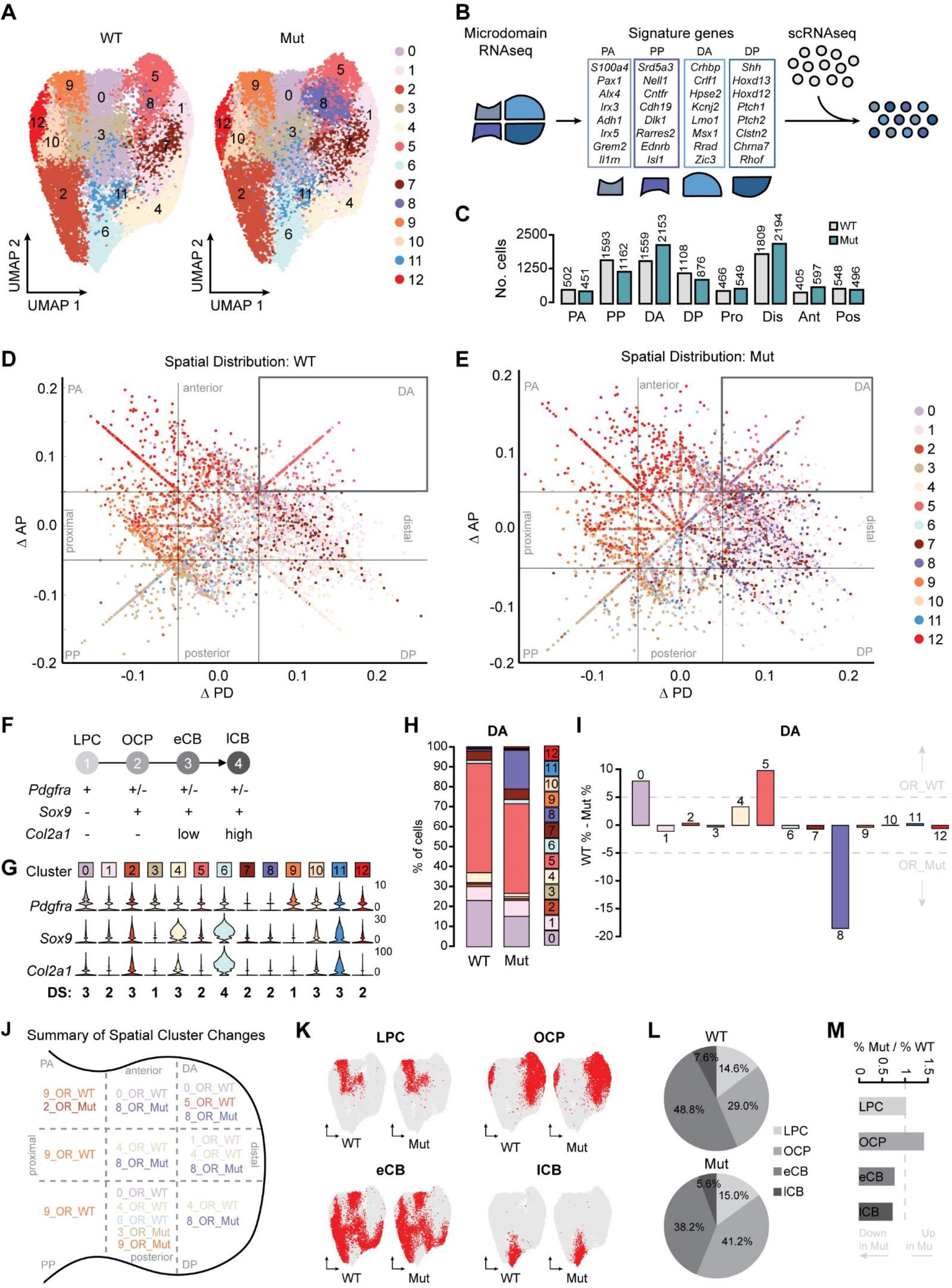
Spatially heterogeneous changes in cell type abundance in mutant forelimb. (A) UMAP of integrated WT and mutant (Mut) endogenous limb mesenchymal cells split by their genotypic origin. (B) Schematic showing workflow for deciphering scRNAseq spatial distribution using microdomain RNAseq data. (C) Bar chart showing number of ELM cells from scRNAseq binned into each microdomain. (D-E) Spatial distribution of all (D) WT and (E) Mut cells. (F) Schematic showing classification scheme for osteochondroprogenitor differentiation state. (G) Violin plot showing expression of *Pdgfra*, *Sox9*, and *Col2a1* for each cluster with classified differentiation state (DS) beneath. (H-I) Bar chart showing the (H) raw percent composition of each cluster within the DA microdomain for WT and mutant with (I) percentage change for each cluster in the mutant. Dashed line in (I) shows cut-off for over-representation (OR). (J) Overall summary of cell type abundance changes in each microdomain of the mutant forelimb. (K) UMAPs showing LPCs, OCPs, eCBs, and lCBs in the WT and Mut datasets. (L) Pie-chart showing cell type composition of the WT and Mut forelimb based on DS. (M) Bar chart showing change in cell type abundance based on DS in the Mut forelimb.

Having approximated each cell’s position and differentiation state, we next aimed to merge these two features in order to determine the relative composition of cells within each WT and mutant microdomain. For this, we quantified the percentage of cells from each cluster that were binned into a specific microdomain and subsequently determined whether this distribution was maintained in the mutant. For example, WT DA cells were predominantly from clusters 0 (early-stage CBs) and 5 (OCPs) (Fig. 5H). Using a 5% cut-off, we found that the mutant DA showed a decrease in the number of cells from both of these clusters with a concomitant increase in the number of cells from cluster 8 (OCPs), which were almost entirely absent in the WT (Fig. 5H-I). Further analysis of the top transcriptional markers of cluster 8 unsurprisingly revealed that this population of cells were undergoing p53-mediated apoptosis, given their expression of *Pmaip1, Ccng1, Cdkn2a*, and *Phlda3* (Supp. Fig. 8A). The WT cells of this cluster expressed *Pmaip1*, but did not express the other markers (Supp. Fig. 8A). This agrees with our discovery through our *spatialPlot* analysis that *Pmaip1* is a novel marker of the DA microdomain, which we confirmed through WISH (Fig. 3A; Supp. Fig. 8B).

Ultimately, performing this analysis for all microdomains revealed regionally-biased cell type alterations in the mutant forelimb (Fig. 5J). For example, cells from cluster 8 (dying OCPs) were over-represented in the mutant center, anterior, DA, distal, and DP microdomains, whereas cells from cluster 4 (late-stage CBs) were over-represented in the WT (and therefore depleted in the Mut) center, distal, DP, and posterior microdomains (Fig. 5J). From this approach, we noticed that most clusters over-represented in the WT forelimb reflected cells of later chondrogenic differentiation (Fig. 5G, J). Thus, we determined the relative composition of cells along the chondrogenic differentiation state, regardless of cluster identity, for both the WT and mutant forelimb (Fig. 5K). Whereas the number of LPCs was approximately equivalent, we found a dramatic increase in the number of OCPs with a concomitant decrease in the number of early- and late-stage chondroblasts in the mutant forelimb, which we interpreted as cellular developmental delay (Fig. 5L-M). Referring back to the spatial location of these cells, we found that developmental delay of osteochondroprogenitors was most likely due to cell abundance changes occurring distally and posteriorly, which aligns with the biased distribution of cell death (Fig. 2A, 5J). Together, the integration of our analyses has allowed us to decipher the impact of molecular heterogeneity on cellular heterogeneity.

### Development of pre-chondrocytes, myoblasts, and axonal branches is delayed in the U11-null forelimb

Regionally distinct splicing alterations are directly connected to discordant gene expression patterns in the U11-null limb, which together could compromise proper limb formation. Indeed, we discovered developmental delay of limb progenitor cells in regard to their ability to progress through the chondrogenesis differentiation program (Fig. 5L-M). Thus, we sought to determine how these molecular and cellular deficits might culminate in perturbed development for three major systems of the limb: the endogenous limb skeletal system and the non-endogenous, invasive muscular and nervous systems^43^. For this, we performed IF for Sox9, Myod1, and Neurofilament-M, which mark chondroprogenitors, myoblasts, and migrating axons, respectively^58,59^, on serial sections collected from WT and mutant E10.5, E11.5, and E12.5 forelimb buds (Fig. 6; Supp. Fig. 9; Supp. Fig. 10; Supp. Fig. 11). We then compiled the staining patterns across each section collected from each individual limb to render a 3-dimensionsal reconstruction for each system (Fig. 6; Supp. Fig. 9; Supp. Fig. 10; Supp. Fig. 11).

**Figure 6.**
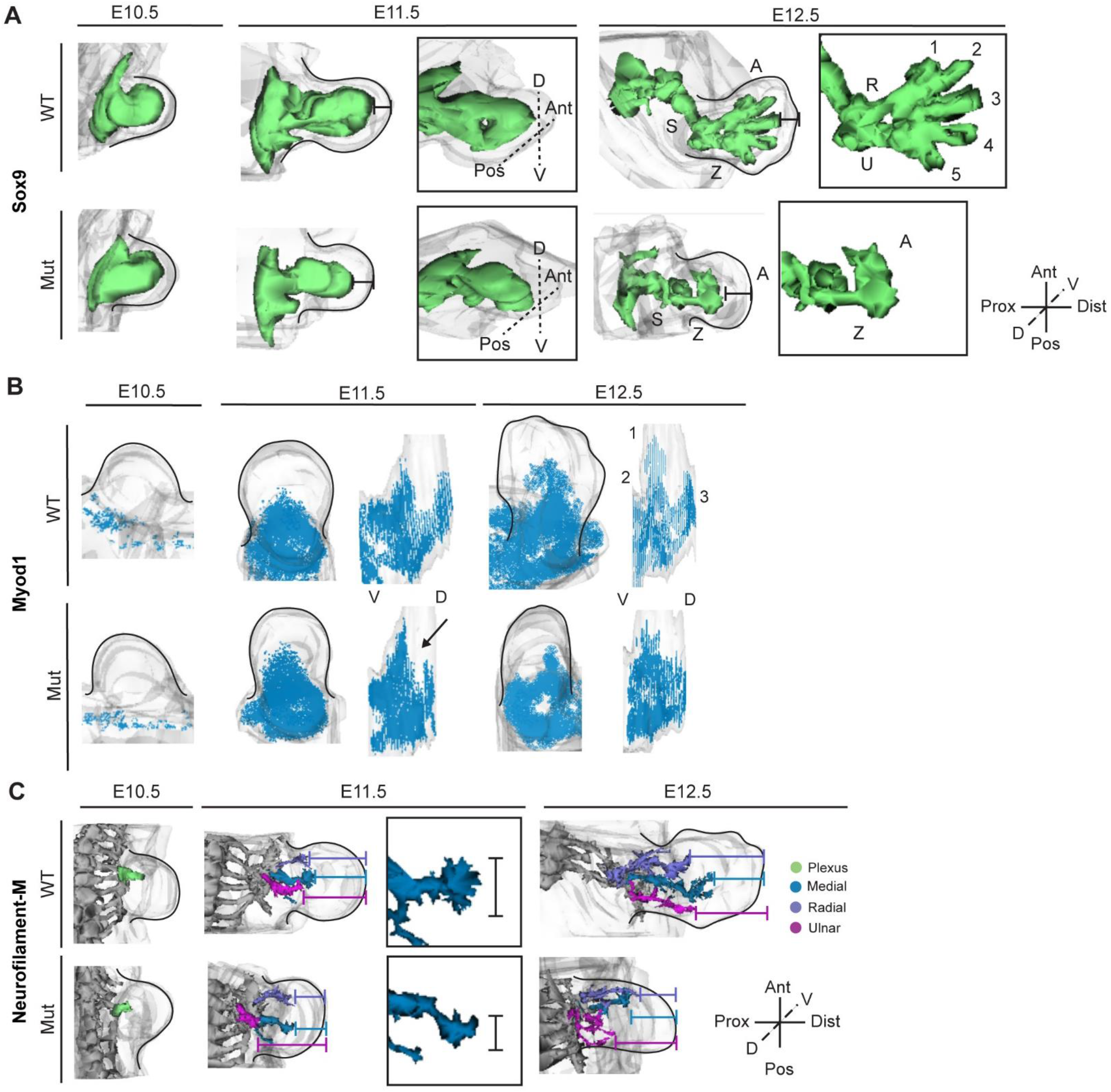
U11 loss results in aberrant chondrogenesis, myogenesis, and axonogenesis. (A-C) Three-dimensionally reconstructed immunofluorescence staining patterns for Sox9 (A), Myod1 (B), and Neurofilament-M (C) in WT and mutant (Mut) forelimbs at E10.5, E11.5, and E12.5. Arrowhead in B shows lack of Mut dorsal muscle mass. Reconstructions are scaled between equivalent WT and Mut pairs, but not between time points or target system. S=stylopod, Z=zeugopod, A=autopod, R=radius, U=ulna, 1-5=digits (A), 1-3=muscle masses (B), Prox=proximal, Dist=distal, D=dorsal, V=ventral, Ant=anterior, Pos=posterior.

We first assessed the patterning and differentiation integrity of the endogenous skeletal program in the U11-null forelimb. Consistent with previous findings, 3D reconstructions of the WT Sox9 profile showed that it is expressed throughout the limbs and somites at E10.5^60^ (Fig. 6A; Supp. Fig. 9). By E11.5, the WT Sox9 expression expands proximo-distally with increased limb outgrowth, and bifurcation of the of the zeugopod can be seen (3/3 reconstructions) (Fig. 6A; Supp. Fig. 9). At this time, the mutant limb shows decreased expansion of Sox9 expression in the distal direction, and simultaneously fails to undergo zeugopod bifurcation (2/3 reconstructions) (Fig. 6A; Supp. Fig. 9). At E12.5, the WT shows a distinct scapula, stylopod, bifurcated zeugopod, and autopod with five digits (Fig. 6A; Supp. Fig. 9). In contrast, the mutant forelimb contains a dysplastic scapula, rudimentary stylopod, a single zeugopodial element (whose posterior location suggests ulnar identity), and an autopodial anlage that lacks any distinct digital condensations (2/3 reconstructions) (Fig. 6A; Supp. Fig. 9). In addition, the length between the distal-most Sox9 expression and the apical ectodermal ridge is greater in the mutant than in the WT, suggesting that distal limb progenitor cells fail to differentiate into chondroprogenitors (Fig. 6A; Supp. Fig. 9).

The migration of muscle progenitors and axons depends crucially on proper patterning and differentiation of the skeletal precursors^47,61–63^. Given that inhibition of the minor spliceosome disrupts endogenous chondroprogenitor development, we hypothesized that migration of both systems would be impaired. We first tested this hypothesis by assessing the spatiotemporal distribution of myoblasts in the WT and U11-null forelimbs (Fig. 6B; Supp. Fig. 10). In agreement with previous publications, we found that Myod1+ muscle progenitors are restricted to the dermomyotome in the WT at E10.5^64^, which was also observed in the mutant (Fig. 6B; Supp. Fig. 10). At E11.5, the WT shows substantial migration of muscle precursors into the proximal portion of the limb bud, which form into two distinct dorsal and ventral muscle masses (Fig. 6B; Supp. Fig. 10). Notably, the mutant forelimb does show appropriate expansion of Myod1+ cells into the proximal limb field; however, we observed a dramatic decrease in the distal expansion of the dorsal muscle mass (Fig. 6B; Supp. Fig. 10). By E12.5, the WT limb shows increased distal migration of myoblasts, which have begun separating into three distinct primordial muscle blocks (Fig. 6B; Supp. Fig. 10). In contrast, muscle patterning along all three axes of limb development appears disorganized and no clear muscle blocks can be identified in the E12.5 mutant forelimb (Fig. 6B; Supp. Fig. 10).

We further interrogated our hypothesis by determining the spatiotemporal distribution of migrating axons in the WT and mutant forelimb (Fig. 6C; Supp. Fig. 11). Axonal projection number, location, and morphology within the developing limb have previously been defined^47,59^. In line with previous reports, we found that the E10.5 WT forelimb contains a plexus of axons in the proximal limb^47,59^ (Fig. 6C; Supp. Fig. 11). This axonal plexus was observed in the E10.5 mutant forelimb at the appropriate position, although it appeared hypoplastic (Fig. 6C; Supp. Fig. 11). By E11.5, the WT forelimb contains 3 main branches invading the proximal limb field, including the radial, ulnar, and medial projections (Fig. 6C; Supp. Fig. 11). Although all three axonal projections could be detected in the E11.5 mutant forelimb, we found that the ulnar nerve showed failed proximo-distal elongation and the medial nerve showed a decrease in density (Fig. 6C; Supp. Fig. 11). At E12.5, all three branches become further elaborated, with the radial, medial, and ulnar nerves greatly increasing their distal migration into the WT limb (Fig. 6C; Supp. Fig. 11). In contrast, in the U11-null forelimb, the three branches become increasingly disordered, with the medial and radial nerves showing anterior displacement and the ulnar nerve appearing coiled rather than being linear (Fig. 6C; Supp. Fig. 11). Taken together, as the mutant forelimb develops, all three systems show increasingly aberrant patterning with simultaneously perturbed distalization (Fig. 6; Supp. Fig. 9; Supp. Fig. 10; Supp. Fig. 11).

## Discussion

Growth and patterning of the developing limb is an orchestrated process built from multiple morphogen gradients that influence cell proliferation and fate^20,35^. Evolutionary-mediated and experimentally-induced alterations to these networks have revealed that the developing limb bud is plastic as it can produce vastly diverse bauplans^1,2^. However, a macroscopic assessment of these structures suggests that the developmental robustness of the limb is preferentially biased for proximo-distal segmentation, as most species display a tripartite limb containing a stylopod, zeugopod, and autopod independent of changes in the number of skeletal elements within each of these units^1,2^. The molecular processes governing this phenomenon remain underexplored.

We reasoned that human disorders wherein individuals show micromelia (an abnormal reduction in limb size) without alteration in their limb pattern, such as MOPD1^12,13^, RS^15^, and LWS^14^, offer a unique opportunity to explore the developmental robustness of the limb at a molecular level. Thus, we used our U11 cKO mouse^65^ to inhibit the minor spliceosome in the developing limbs, which we have previously shown to result in similar yet more severe micromelia as observed in minor spliceosomopathies^11,16^. Using this system, we discovered that minor spliceosome inhibition results in transcriptomic stalling, as genes that should be downregulated across time were unchanged in the mutant forelimb (Fig. 1A-H). We captured transcriptional stalling between E10.5 and E11.5, which is when cell cycle defects and cell death are precipitating in the mutant forelimb^11^ (Fig. 2A). Thus, it is unclear whether transcriptional stalling contributes to or is a result of these cellular deficits. In addition, it remains unexplored whether transcriptional stalling is a general outcome of global LPC depletion or specific to minor spliceosome inhibition. Multiple MIGs regulate chromatin architecture and transcription^24^; thus, their mis-splicing in our system may uniquely drive transcriptional stalling. However, it is equally likely that progression of the limb transcriptional program is tied to appropriate establishment of LPC proliferation and morphogen signaling. In this case, transcriptional stalling might reflect a holding period for the limb to buffer developmental insults. Favoring the latter, transcriptional stalling in the mutant forelimb affected numerous *Hox* genes, which are non-MIGs crucial for proper limb development^19^ (Fig. 1G, H). Using *Hoxa11* as an exemplar, we found this molecular perturbation to be regionally specific and was sufficient to disrupt its encoded protein function (Fig. 1I-L). Interestingly, one of the over-bound peaks in the *Hoxa11* dataset corresponds to the *Prrx1* promoter. For this, there are two possibilities: 1) our use of the *Prrx1*-Cre^17^ mouse line provides three promoters for Hoxa11 to bind; or 2) over-bound Hoxa11 at the *Prrx1* promoter is biologically relevant. We favor the latter possibility because *Prrx1* is a marker of early LPCs^17^, which agrees with transcriptional stalling of the E11.5 mutant forelimb. In contrast, under-bound promoters included *Noc2l*, a histone deacetylase inhibitor known to regulate p53-mediated apoptosis^23^. Noc2l prevents p53 from binding to its target promoters, effectively inhibiting apoptosis^23^. This suggests that reduced transcriptional activation of Noc2l could allow p53 to activate its downstream effectors, an idea supported by increased levels of apoptosis in the mutant forelimb^11^ (Fig. 2A). Investigations of temporal transcriptional dynamics in other models of limb development defects is thus warranted to determine if transcriptional stalling is a conserved phenomenon.

The developing limb is intricately organized with spatially distinct cell populations governing growth, patterning, and differentiation^20,35^. We have previously found that cellular consequences of minor spliceosome inhibition preferentially affect distal posterior LPCs^11^ (Fig. 2A). Thus, we devised our microdomain method to spatially profile the molecular changes occurring within the mutant forelimb (Fig. 2B). This approach uncovered intra-tissue specific mis-splicing of MIGs in the mutant forelimb, which importantly expands our appreciation for splicing heterogeneity beyond the previously known inter-tissue specific differences^24^ (Fig. 2B-L). This data can be leveraged toward resolving why distal posterior LPCs are more susceptible to apoptosis upon U11 loss^11^, as rapid cell death might be facilitated by unique mis-splicing events in this microdomain. In turn, these findings can be broadly extended toward resolving the paradoxical, tissue-specific pathology of spliceosomopathies^66,67^.

Here, our spatial transcriptomics analysis suggested a failure of the mutant forelimb to separate its distal patterning program from its proximal differentiation program, which is likely an extension of transcriptional stalling (Fig. 3A-G). As transcriptional profiles reflect cell identities, this perturbation led us to suspect that the progression of LPCs along the chondroprogenitor trajectory was disrupted. Indeed, through single cell RNAseq, we found that the mutant forelimb displayed developmental delay of chondroprogenitors as reflected by an accumulation of OCPs and a subsequent depletion of early- and late-stage CBs (Fig. 5L-M). Importantly, integration of our spatial transcriptomic data with our single cell RNAseq data allowed us to map the relative spatial position of each cell, revealing that most cell type abundance alterations were occurring distally and posteriorly (Fig. 5J). It remains unclear whether this is due to cell death cues, unique mis-splicing events, or other molecular perturbations in this region serving to repress the developmental trajectory of OCPs.

Altered regional transcription and cell-type abundances prompted us to investigate the 3D patterning of multiple systems of the developing mutant forelimb. We discovered that the micromelic phenotype of the mutant forelimb^11^ precipitates as early as E12.5, where developmental delay of chondroprogenitors severely perturbs patterning of the anterior zeugopod and the autopod (Fig. 6A). Notably, the absence of the anterior zeugopod and the presence of an autopodial anlage emphasizes the aforementioned biological phenomenon; upon depletion of LPCs, the mutant forelimb displays a developmental bias towards proximo-distal segmentation in lieu of antero-posterior bifurcation (Fig. 6A). Moreover, the lack of an anterior zeugopodial element was perplexing as the most notable cell type abundance changes affecting chondroblast production were distal posteriorly concentrated (Fig. 5J). How the mutant forelimb develops a posterior zeugopod upon massive apoptosis and chondroblast depletion remains unclear (Fig. 2A, 5J).

As the endogenous limb mesenchyme must coordinate the migration and patterning of muscle and nervous tissue during development^43,59^, we suspected that the ultimate culmination of developmental delay would be disrupted development of these non-endogenous systems. Indeed, we found perturbed patterning of both myoblasts and axonal branches (Fig. 6B, C). One of the primary limitations for our microdomain analysis is that it omits differences along the dorsoventral axis, which are equally crucial for limb development and patterning^68^ (Fig. 2B). This deficit is particularly challenging to overcome in a reproducible manner given how narrow the dorsoventral axis is at the embryonic stages interrogated. Devising a method to surmount this limitation would prove valuable, as our analyses revealed a major deficit in migrating myoblasts to form dorsal muscle masses (Fig. 6B; Supp. Fig. 10). As most of these cells do not express *Prrx1*, their perturbed migration and patterning is a secondary outcome of a failure of the U11-null limb to establish an appropriate spatial cell-signaling program^17^. Interestingly, the substantial defect in distal outgrowth for the ulnar nerve, as well as the anterior displacement for the medial and radial nerves, suggests that cell death may serve as a repulsive cue for axonal migration (Fig. 2A, 6C; Supp. Fig. 11). Future experiments leveraging ablation of p53 (to block LPC apoptosis) in the U11-null forelimb would help resolve primary phenotypes occurring from mis-splicing of MIGs and secondary phenotypes resulting from cell death^69^. Nonetheless, our systems analysis reveals that the minor spliceosome is required not only for the proper development of endogenous limb progenitors, but also to generate the molecular framework from which the entire limb will be constructed (Fig. 6; Supp. Fig. 9; Supp. Fig. 10; Supp. Fig. 11).

Altogether, our data reveals that upon minor spliceosome inhibition, the developing limb undergoes transcriptomic stalling leading to cellular developmental delay. We posit that this mechanism, which may be a general phenomenon triggered by LPC depletion, affords the limb the opportunity to pause its development in order to buffer organogenetic insults. When left unresolved, as is the case for our system, these deficits culminate in failed formation for both endogenous and non-endogenous systems.

## Acknowledgements

We thank Dr. Bo Reese from the University of Connecticut’s Center for Genome Innovation for assistance with RNA/DNA sequencing; Dr. Jeremy Balsbaugh and Dr. Jen Liddle from the University of Connecticut’s Proteomics & Metabolomics Facility for assistance with mass spectrometry sample preparation, data acquisition, and computational analysis; and Dr. Ion Mandoiu from the University of Connecticut’s Computer Science and Engineering Department for the establishment of bioinformatic platforms. The authors would also like to acknowledge the NIH S10 High-End Instrumentation Award 1S10-OD028445-01A1, which supported this work by providing funds to acquire the Orbitrap Eclipse Tribrid mass spectrometer housed in the University of Connecticut Proteomics & Metabolomics Facility. Support for this work was provided by funding from NSF GRFP 2018257410 to K.D.D. and R01 NS102538 to R.N.K.

## Author Contributions

Conceptualization, K.D.D., S.M.S., R.N.K.; Methodology, K.D.D., S.M.S., R.N.K.; Software, K.D.D., S.M.S.; Validation, K.D.D., S.M.S., K.O.A., T.D.L., K.N.G.; Formal Analysis, K.D.D., S.M.S., R.N.K.; Investigation, K.D.D., S.M.S., K.O.A., T.D.L., K.N.G., R.N.K.; Resources, R.N.K.; Data Curation, K.D.D., S.M.S., R.N.K.; Writing – Original Draft, K.D.D., S.M.S., R.N.K.; Writing – Review & Editing, K.D.D., S.M.S., R.N.K.; Visualization, K.D.D., S.M.S., R.N.K.; Supervision, R.N.K.; Project Administration, R.N.K.; Funding Acquisition, K.D.D., R.N.K.

## Declaration of Interests

The authors declare no competing or financial interests.

## Methods

### Animal husbandry

Mouse care and experimental use was carried out along guidelines established by the University of Connecticut’s Institutional Animal Care and Use Committee. This study employed the *Rnu11* conditional knockout mouse generated by Baumgartner et al.^65^ crossed with *Prrx1-Cre*^17^, which was originally reported in Drake et al.^11^ Wild-type (WT) animals were either *Rnu11*^WT/Flx^::*Prrx1-Cre*^-^ or *Rnu11*^Flx/Flx^::*Prrx1-Cre*^-^. Mutant animals were *Rnu11*^Flx/Flx^::*Prrx1-Cre*^+^. Male and female embryos were used equally. Embryonic staging was carried out such that E0.5 was considered to be noon after a morning vaginal plug was observed. Each mutant embryo used in this study had a comparably staged littermate control.

### Bioinformatics analysis & data accessibility

#### Total RNAseq

Total RNAseq was performed on E10.5 and E11.5 WT and mutant forelimb and hindlimb (N=3 for each sample). RNA extraction was done using TRIzol (Thermo Fisher Scientific, 15596018) per the manufacturer’s instructions. RNA sample preparation and sequencing were executed by the University of Connecticut’s Center for Genome Innovation. Library prep was done through Illumina TruSeq Stranded Total RNA Library Sample Prep Kit (RS-122-2201) with RiboZero for ribosomal RNA depletion. Sequencing was performed using Illumina NextSeq 500. Reads were mapped to mm10 using Hisat2^70^. Gene expression calls were determined through IsoEM2^34^. Differential gene expression was determined using IsoDE2^34^. DAVID was employed for functional enrichment analysis of gene sets with significance determined by Benjamini-Hochberg adjusted *P*-value<0.05. Bulk RNAseq data used in this manuscript can be accessed through NCBI’s Gene Expression Omnibus Series accession number GSE146424.

#### PCA

Principal component analysis was performed on log-normalized TPM values through ClustVis^71^ using default settings.

#### CUT&RUN

Embryos were harvested at E11.5 and N=3 WT and mutant forelimbs (pooled left and right) were dissociated into single cell suspensions using collagenase (Sigma C9697; 50mg/mL) at 37°C for 15 minutes followed by gentle pipette mixing. Samples were processed using the EpiCypher CUTANA™ CUT&RUN Kit (SKU: 14-1048) per the manufacturer’s instructions with 0.5 μg/100 μl anti-Hoxa11 (Sigma-Aldrich SAB1304728) or anti-IgG (CUTANA™ SKU: 13-0042). Libraries were prepped using NEBNext Ultra™ II DNA Library Prep Kit for Illumina per manufacturer’s instructions. Sequencing was conducted through the University of Connecticut’s Institute for Genome Innovation using Illumina NovaSeq SP 200 cycle for paired-end, 100 bp reads at 8-10 million read depth per sample. For computational analysis, adapters and bad-quality bases were removed with Cutadapt 3.5; reads with poor mapping quality (below 30 or unpaired) were removed with fastx; reads were mapped to mm10 using Bowtie2 (-I 0 -X 1000 –dovetail –very-sensitive); PCR duplicates were removed with Picard 2.9.2; coverage was identified with MACS2 (--format BED –keep-dup all –bdg –nomodel –extsize 200 – shift −100); Hoxa11 binding heatmaps were generated using deepTools plotHeatmap (--binSize 20 – normalizeUsing BPM –scaleFactorsMethod None –smoothLength 60 –extendReads 150 –centerReads) and were normalized using appropriate sample control IgG binding profiles; MOTIF enrichment was identified using HOMER 4.10; and differential binding analysis was conducted using DiffBind 2.14 using default conditions via DESeq2. The data presented in this manuscript is currently in process of submission to NCBI’s Gene Expression Omnibus. Reviewer access token will be granted upon request.

#### Microdomain RNAseq

Total RNAseq was performed on N=3 E11.5 WT and mutant proximal anterior, proximal posterior, distal anterior, and distal posterior forelimb microdomains, of which left and right microdomains were pooled for each individual N. TRIzol-based RNA extraction was employed using instructions provided by the manufacturer. Through the University of Connecticut’s Center for Genome Innovation, RNA samples underwent library preparation using the Illumina TruSeq Stranded Total RNA Library Sample Prep Kit, were ribo-depleted using RiboZero, and were sequenced using Illumina NovaSeq 6000. Reads were mapped to mm10 using Hisat2^70^. Differential gene expression was determined using IsoEM2 and IsoDE2^34^. Minor intron retention and alternative splicing analyses were conducted as reported in Olthof et al.^24^ DAVID was used to identify biological pathway enrichment, with only KEGG pathways and GO terms having Benjamini-Hochberg-adjusted *P*-value<0.05 included for analysis. STRING was used to determine gene connectivity networks. The data presented in this manuscript is currently in process of submission to NCBI’s Gene Expression Omnibus. Reviewer access token will be granted upon request.

#### Untargeted Protein Identification and Label-free Quantification via Tandem Mass Spectrometry

Submitted samples were dried to completion and resuspended in 23 μL of lysis buffer (50 mM ammonium bicarbonate, 5% SDS) prior to preparation with S-Trap columns (ProtiFi, LLC) using a modified commercial protocol. Samples were first reduced with 120 mM dithiothreitol for 1 hr at room temperature, then alkylated with 500 mM iodoacetamide for 30 min in the dark at room temperature, then acidified to a final concentration of 2.5% phosphoric acid, and finally diluted 6-fold with 0.1 M triethylammonium bicarbonate (TEAB) in 90% methanol (MeOH). Diluted samples were loaded on a Micro S-Trap (ProtiFi part #02-micro-80), washed per commercial protocol with 0.1 M TEAB in 90% MeOH, and incubated with sequencing grade, modified porcine trypsin (Promega #V5113) at 1:10 (w/w) enzyme:sample overnight at 25°C. Peptides were eluted with successive washes of 50 mM TEAB, 0.2% formic acid in H-2O, and 0.2% formic acid in acetonitrile, pooled, and dried. Peptides were then desalted by reverse-phase chromatography using Pierce peptide desalting spin columns (Thermo Fisher part #89852). Final resuspension was done with Solvent A (0.1% formic acid in Fisher Optima LC/MS grade water) and samples were quantified by A-280 absorbance. Total injection amount by mass was normalized across all samples.

Samples were analyzed using a Thermo Scientific Ultimate 3000 RSLCnano ultra-high performance liquid chromatography (UPLC) system coupled to a high-resolution Thermo Scientific Eclipse Tribrid Orbitrap mass spectrometer. Each sample was injected onto a nanoEase M/Z Peptide BEH C18 column (1.7 μm, 75 μm x 250 mm, Waters Corporation) and separated by reversed-phase UPLC using a gradient of 4-30% Solvent B (0.1% formic acid in Fisher Optima LC/MS grade acetonitrile) over a 150-min gradient at 300 nL/min flow, followed by a ramp of 30-90% Solvent B over 30 min, for a 180-min total gradient. Peptides were eluted directly into the Eclipse using positive mode nanoflow electrospray ionization with a capillary voltage of 2200 V. MS1 scans were acquired at 120,000 resolution, with an AGC target of 4e5, maximum ion time set to Auto, RF lens setting of 30%, and a scan range of 300-1800 m/z. Data-dependent MS2 scans were acquired in the ion trap using a Rapid scan rate, for any ions over 5.0e3 intensity threshold, with AGC target set to Standard, maximum ion time set to Auto, mass range set to Normal, scan range set to Auto, isolation window set to 1.6 m/z, and a fixed cycle time of 3 sec. CID fragmentation was applied with 35% energy and an activation time of 10 ms. Dynamic exclusion was set to exclude for 30 sec after 1 observation, and charge states of 2-8 were included.

Peptides were identified using MaxQuant software (v1.6.10.43) and its embedded Andromeda search engine, and quantified using label-free quantification^72^. The raw data were searched against both the complete UniProt *Mus Musculus* reference proteome (identifier UP000000589, accessed 01/12/2022) and the MaxQuant contaminants database. Minimum peptide length was set to 5 residues. Variable modifications allowed oxidation of Met, acetylation of protein N-termini, deamidation of Asn/Gln, and peptide N-terminal Gln to pyroGlu conversion. Carbamidomethylation of Cys was set as a fixed modification. Protease specificity was set to trypsin/P with a maximum of 2 missed cleavages. All results were filtered to a 1% false discovery rate at the peptide and protein levels using the target-decoy approach; all other parameters were kept at default values. MaxQuant output files were imported into Scaffold (v5.1.2, Proteome Software, Inc.) for data visualization and subsequent analyses.

#### Differential Protein Expression Analysis

Average Precursor Identity (API) was used for differential protein expression analysis. Samples containing replicates with missing protein expression values were imputed by averaging the API values of all three replicates. A minimum of two replicates with API values > 1e5 were required for a protein to be considered expressed in a sample. Student’s two-tailed T-test with Benjamini-Hochberg adjustment was run on log-transformed API values between appropriate WT and mutant sample pairs to determine differential protein expression with *P*<0.05 used as cut-off for significance.

#### spatialPlot

Regional gene expression was determined for each N by averaging proximal (PA, PP), distal (DA, DP), anterior (PA, DA), or posterior (PP, DP) microdomain transcript per million (TPM) values. Subsequently, log 2 fold change (log2FC) values were determined for each axis (distal/proximal; anterior/posterior) and plotted, with log2FC ≥ 1 or ≤ −1 used as cut-off for enrichment in each domain. In addition, microdomain gene expression values were compared all-to-all using ANOVA with *post-hoc* Tukey’s test and Benjamini-Hochberg *P*-value adjustment. The following criteria were used to define significant enrichment for each region: PA enriched = PA TPM significantly greater than PP, DA, and DP TPM; PP enriched = PP TPM significantly greater than PA, DA, and DP TPM; DA enriched = DA TPM significantly greater than PA, PP, and DP TPM; DP enriched = DP TPM significantly greater than PA, PP, and DA TPM; proximally enriched = PA TPM significantly greater than DA and DP TPM, and PP TPM significantly greater than DA and DP TPM; distally enriched = DA TPM significantly greater than PA and PP TPM, and DP TPM significantly greater than PA and PP TPM; anteriorly enriched = PA TPM significantly greater than PP and DP TPM, and DA TPM significantly greater than PP and DP TPM; posteriorly enriched = PP TPM significantly greater than PA and DA TPM, and DP TPM significantly greater than PA and DA TPM. A gene had to meet both the log2FC threshold and statistical significance by ANOVA to be determined spatially restricted in *spatialPlot* (note: muted color dots in Fig. 3A-B show genes that meet the log2FC thresholding but were not significantly enriched by ANOVA).

#### Single Cell RNAseq

Forelimbs dissected from N=6 E11.5 WT and mutant embryos were pooled into two batches, with each batch containing one limb from each embryo (i.e., WT 1 left forelimb in WT batch 1, WT 1 right forelimb in WT batch 2). Left and right forelimbs were intermixed for each batch. Single cell suspensions were generated using collagenase (Sigma C9697; 50mg/mL) at 37°C for 15 minutes followed by gentle pipette mixing. Single cell capture and library preparation was carried out through the University of Connecticut’s Center for Genome Innovation. Each sample was processed with 10x Genomics NextGem v3.1 kit, per the manufacturer’s instructions. Samples were sequenced on a NovaSeq SP100 cycle v1.5. Resulting .bcl files were converted to fastq files and mapping was performed using cellranger mkfastq and cellranger count, respectively. WT and mutant batches were then merged using cellranger aggr with default settings. Output data was analyzed using Seurat^73^ with filtering criteria as reported by Kelly et al.^56^, wherein cells expressing less than 200 genes, more than 6000 genes, or containing >10% mitochondrial reads were removed from analysis. Gene expression counts were then log normalized and the top 1000 variable features were determined using vst. Principal component analysis (PCA) was run on the top 20 significant dimensions and cluster identification was generated through UMAP using default settings (resolution=0.5). The top gene expression markers for each cluster were extracted using Seurat’s FindAllMarkers^73^ with default parameters. Cluster labeling was facilitated by using markers reported in Kelly et al.^56^ and Reinhardt et al.^57^ Endogenous limb mesenchyme cell populations were regressed for cell cycle score post-integration.

#### Integration of Spatial Transcriptomics with Single Cell RNAseq

Signature genes were determined as described in Supplementary Figure 6. The top 8 signature genes for each microdomain were then used for input lists into Ucell’s_addModuleScore function to add a similarity score for each microdomain to every cell. Regional module scores were then generated by averaging the two corresponding microdomain scores for said region (i.e., average of DA & PA = anterior score). Subsequently, the difference between the distal and proximal scores as well as between the anterior and posterior scores was calculated. A difference threshold of 0.05 (a value slightly larger than the standard deviation of each difference dataset) was used to define the 9 spatial bins (Fig. 5D-E). Validation of this binning method is shown in Supplementary Figure 7.

Percent composition of each cluster for each microdomain was calculated by dividing the number of cells binned in a specific spatial location and cluster by the total number of cells binned in said spatial location (ex: the number of cluster 2 cells in DA divided by the total number of DA-binned cells).

### RT-PCR

cDNA was synthesized from RNA extracted from N=3 WT and mutant microdomains, which was then pooled for RT-PCR. Forward and reverse primers were designed two exons upstream and two exons downstream the minor intron in the MIG of interest, respectively (sequences in Table 1). Differentially spliced transcripts identified in our RNAseq analysis were confirmed by their predicted molecular weight through agarose gel electrophoresis.

### Immunofluorescence and 3D reconstructions

WT and U11-null forelimbs were collected at E10.5, E11.5, and E12.5 and prepared for mounting and sectioning by incubating in 4% PFA/PBS overnight at 4°C. The tissue was then washed twice with PBS and incubated in 30% sucrose/PBS overnight at 4°C. Following a final overnight incubation with a 1:1 ration of OCT and 30% sucrose/PBS, limbs were mounted in OCT and stored at −80°C until sectioning. Thirty minutes prior to sectioning, the tissue was stored at −20°C. Serial cryosections of 25 μm thickness were collected for N=1 WT and U11-null limb bud per slide set. Immunofluorescence was performed as previously described^11^. Primary antibodies were diluted to 1:500 (mouse anti-Myod1, BD pharmaceuticals, 554130; rabbit anti-Sox9, Millipore, AB5535), 1:300 (rabbit anti-cleaved caspase 3, Cell Signaling, #9665S), or 1:200 (rabbit anti-Neurofilament-M, Millipore, AB1987). Both left and right limbs were sectioned and reconstructed for each timepoint and each antibody. Serial sections were then imaged on the Keyence BZ-X710 at 10x with automatic settings for signal intensity and image stitching. Serial images were then imported into Reconstruct into individual series for WT and U11-null images. Images were calibrated by drawing a single trace the length of a 50 μm scale bar. These images were then manually aligned to one another and signal for either CC3 (N=1), Sox9 (N=3), Myod1 (N=3), or Neurofilament-M (N=2) was traced in addition to the outline of the limb bud. 3D objects were then reconstructed by generating a boissonnat surface from the individual traces using the original settings from Reconstruct^74^.

### Whole mount *in situ* hybridization

Whole mount *in situ* hybridization was performed as described in Drake et al.^11^ The following sequences were used as primers to generate probes: *Hoxa11* – forward 5’ – GGTTCAGATCTCCGTGGTTAAG – 3’; reverse 5’ – GGCTCTTAGAAGATTGCCAGAA – 3’; *Pmaip1* – forward 5’ – CAGGGTTCCTTTCCTCTGTGTAGC – 3’; reverse 5’ – TCCCATCACTTCTGGTAGGCATCA – 3’.

## Notes

### Competing Interest Statement

The authors have declared no competing interest.

## References

1. Zuniga, A. Next generation limb development and evolution: old questions, new perspectives. Development 142, 3810–3820 (2015).

2. Petit, F., Sears, K. E. & Ahituv, N. Limb development: a paradigm of gene regulation. Nat Rev Genet 18, 245–258 (2017).

3. Harfe, B. D., McManus, M. T., Mansfield, J. H., Hornstein, E. & Tabin, C. J. The RNaseIII enzyme Dicer is required for morphogenesis but not patterning of the vertebrate limb. Proc Natl Acad Sci U S A 102, 10898–10903 (2005).

4. Singh, J. D. & Singh, S. Skeletal malformations induced by mitomycin C in chick embryos. Acta Orthop Scand 47, 509–514 (1976).

5. Soshnikova, N., Montavon, T., Leleu, M., Galjart, N. & Duboule, D. Functional analysis of CTCF during mammalian limb development. Dev Cell 19, 819–830 (2010).

6. Hom, J. et al. Limb Specific Failure of Proliferation and Translation in the Mesenchyme Leads to Skeletal Defects in Diamond Blackfan Anemia. 2022.01.14.476336 Preprint at https://doi.org/10.1101/2022.01.14.476336 (2022).

7. Sousounis, K. et al. Eya2 promotes cell cycle progression by regulating DNA damage response during vertebrate limb regeneration. eLife 9, e51217 (2020).

8. Ohsugi, K., Gardiner, D. M. & Bryant, S. V. Cell Cycle Length Affects Gene Expression and Pattern Formation in Limbs. Developmental Biology 189, 13–21 (1997).

9. Wolpert, L., Tickle, C. & Sampford, M. The effect of cell killing by x-irradiation on pattern formation in the chick limb. J Embryol Exp Morphol 50, 175–193 (1979).

10. Galloway, J. L., Delgado, I., Ros, M. A. & Tabin, C. J. A reevaluation of X-irradiation-induced phocomelia and proximodistal limb patterning. Nature 460, 400–404 (2009).

11. Drake, K. D., Lemoine, C., Aquino, G. S., Vaeth, A. M. & Kanadia, R. N. Loss of U11 small nuclear RNA in the developing mouse limb results in micromelia. Development 147, dev190967 (2020).

12. He, H. et al. Mutations in U4atac snRNA, a component of the minor spliceosome, in the developmental disorder MOPD I. Science 332, 238–240 (2011).

13. Edery, P. et al. Association of TALS developmental disorder with defect in minor splicing component U4atac snRNA. Science 332, 240–243 (2011).

14. Farach, L. S. et al. The expanding phenotype of RNU4ATAC pathogenic variants to Lowry Wood syndrome. American Journal of Medical Genetics Part A 176, 465–469 (2018).

15. Merico, D. et al. Compound heterozygous mutations in the noncoding RNU4ATAC cause Roifman Syndrome by disrupting minor intron splicing. Nat Commun 6, 8718 (2015).

16. Verma, B., Akinyi, M. V., Norppa, A. J. & Frilander, M. J. Minor spliceosome and disease. Semin Cell Dev Biol 79, 103–112 (2018).

17. Logan, M. et al. Expression of Cre Recombinase in the developing mouse limb bud driven by a Prxl enhancer. Genesis 33, 77–80 (2002).

18. Riddle, R. D., Johnson, R. L., Laufer, E. & Tabin, C. Sonic hedgehog mediates the polarizing activity of the ZPA. Cell 75, 1401–1416 (1993).

19. Zakany, J. & Duboule, D. The role of Hox genes during vertebrate limb development. Current Opinion in Genetics & Development 17, 359–366 (2007).

20. Tabin, C. & Wolpert, L. Rethinking the proximodistal axis of the vertebrate limb in the molecular era. Genes Dev. 21, 1433–1442 (2007).

21. Skene, P. J. & Henikoff, S. An efficient targeted nuclease strategy for high-resolution mapping of DNA binding sites. eLife 6, e21856.

22. McDonel, P., Demmers, J., Tan, D. W. M., Watt, F. & Hendrich, B. D. Sin3a is essential for the genome integrity and viability of pluripotent cells. Dev Biol 363–318, 62–73 (2012).

23. Hublitz, P. et al. NIR is a novel INHAT repressor that modulates the transcriptional activity of p53. Genes Dev. 19, 2912–2924 (2005).

24. Olthof, A. M., Hyatt, K. C. & Kanadia, R. N. Minor intron splicing revisited: identification of new minor intron-containing genes and tissue-dependent retention and alternative splicing of minor introns. BMC Genomics 20, 686 (2019).

25. Olthof, A. M. et al. Disruption of exon-bridging interactions between the minor and major spliceosomes results in alternative splicing around minor introns. Nucleic Acids Res 49, 3524–3545 (2021).

26. Li, F.-Q. et al. BAR Domain-Containing FAM92 Proteins Interact with Chibby1 To Facilitate Ciliogenesis. Mol Cell Biol 36, 2668–2680 (2016).

27. Schrauwen, I. et al. FAM92A Underlies Nonsyndromic Postaxial Polydactyly in Humans and an Abnormal Limb and Digit Skeletal Phenotype in Mice. J Bone Miner Res 34, 375–386 (2019).

28. Datar, K. V., Dreyfuss, G. & Swanson, M. S. The human hnRNP M proteins: identification of a methionine/arginine-rich repeat motif in ribonucleoproteins. Nucleic Acids Res 21, 439–446 (1993).

29. Gattoni, R. et al. The human hnRNP-M proteins: structure and relation with early heat shock-induced splicing arrest and chromosome mapping. Nucleic Acids Res 24, 2535–2542 (1996).

30. Dickinson, M. E. et al. High-throughput discovery of novel developmental phenotypes. Nature 537, 508–514 (2016).

31. Schneider, C. & Tollervey, D. Threading the barrel of the RNA exosome. Trends Biochem Sci 38, 485–493 (2013).

32. Lingaraju, M. et al. To Process or to Decay: A Mechanistic View of the Nuclear RNA Exosome. Cold Spring Harb Symp Quant Biol 84, 155–163 (2019).

33. Shupp, A., Casimiro, M. C. & Pestell, R. G. Biological functions of CDK5 and potential CDK5 targeted clinical treatments. Oncotarget 8, 17373–17382 (2017).

34. Mandric, I. et al. Fast bootstrapping-based estimation of confidence intervals of expression levels and differential expression from RNA-Seq data. Bioinformatics 33, 3302–3304 (2017).

35. Bénazet, J.-D. & Zeller, R. Vertebrate limb development: moving from classical morphogen gradients to an integrated 4-dimensional patterning system. Cold Spring Harb Perspect Biol 1, a001339 (2009).

36. Kuijper, S. et al. Genetics of shoulder girdle formation: roles of Tbx15 and aristaless-like genes. Development 132, 1601–1610 (2005).

37. Probst, S. et al. SHH propagates distal limb bud development by enhancing CYP26B1-mediated retinoic acid clearance via AER-FGF signalling. Development 138, 1913–1923 (2011).

38. Lewandowski, J. P. et al. Spatiotemporal regulation of GLI target genes in the mammalian limb bud. Developmental Biology 406, 92–103 (2015).

39. Galli, A. et al. Distinct roles of Hand2 in initiating polarity and posterior Shh expression during the onset of mouse limb bud development. PLoS Genet 6, e1000901 (2010).

40. Yokouchi, Y., Sasaki, H. & Kuroiwa, A. Homeobox gene expression correlated with the bifurcation process of limb cartilage development. Nature 353, 443–445 (1991).

41. Nagai, T. et al. The Expression of the MouseZic1, Zic2, andZic3Gene Suggests an Essential Role forZicGenes in Body Pattern Formation. Developmental Biology 182, 299–313 (1997).

42. Yokoyama, S. et al. Analysis of transcription factors expressed at the anterior mouse limb bud. PLoS One 12, e0175673 (2017).

43. Martin, P. Tissue patterning in the developing mouse limb. Int J Dev Biol 34, 323–336 (1990).

44. Robertson, E. J. et al. Blimp1 regulates development of the posterior forelimb, caudal pharyngeal arches, heart and sensory vibrissae in mice. Development 134, 4335–4345 (2007).

45. Ratola, A. et al. A Novel Noonan Syndrome RAF1 Mutation: Lethal Course in a Preterm Infant. Pediatr Rep 7, 5955 (2015).

46. Cobb, J., Dierich, A., Huss-Garcia, Y. & Duboule, D. A mouse model for human short-stature syndromes identifies Shox2 as an upstream regulator of Runx2 during long-bone development. Proceedings of the National Academy of Sciences 103, 4511–4515 (2006).

47. Vickerman, L., Neufeld, S. & Cobb, J. Shox2 function couples neural, muscular and skeletal development in the proximal forelimb. Developmental Biology 350, 323–336 (2011).

48. Ye, W. et al. A unique stylopod patterning mechanism by Shox2-controlled osteogenesis. Development 143, 2548–2560 (2016).

49. Neufeld, S. J., Wang, F. & Cobb, J. Genetic Interactions Between Shox2 and Hox Genes During the Regional Growth and Development of the Mouse Limb. Genetics 198, 1117–1126 (2014).

50. Yu, L. et al. Shox2 is required for chondrocyte proliferation and maturation in proximal limb skeleton. Dev Biol 306, 549–559 (2007).

51. Deng, H., Huang, X. & Yuan, L. Molecular genetics of the COL2A1-related disorders. Mutat Res Rev Mutat Res 768, 1–13 (2016).

52. Seghatoleslami, M. R. et al. Differential regulation of COL2A1 expression in developing and mature chondrocytes. Matrix Biol 14, 753–764 (1995).

53. Zhao, Q., Eberspaecher, H., Lefebvre, V. & de Crombrugghe, B. Parallel expression of Sox9 and Col2a1 in cells undergoing chondrogenesis. Developmental Dynamics 209, 377–386 (1997).

54. Mao, J., McGlinn, E., Huang, P., Tabin, C. J. & McMahon, A. P. Fgf-dependent Etv4/5 activity is required for posterior restriction of Sonic hedgehog and promoting outgrowth of the vertebrate limb. Dev Cell 16, 600–606 (2009).

55. Li, C., Scott, D. A., Hatch, E., Tian, X. & Mansour, S. L. Dusp6(Mkp3) is a negative feedback regulator of FGF stimulated ERK signaling during mouse development. Development 134, 167–176 (2007).

56. Kelly, N. H., Huynh, N. P. T. & Guilak, F. Single cell RNA-sequencing reveals cellular heterogeneity and trajectories of lineage specification during murine embryonic limb development. Matrix Biol 89, 1–10 (2020).

57. Reinhardt, R. et al. Molecular signatures identify immature mesenchymal progenitors in early mouse limb buds that respond differentially to morphogen signaling. Development 146, dev173328 (2019).

58. Akiyama, H., Chaboissier, M.-C., Martin, J. F., Schedl, A. & de Crombrugghe, B. The transcription factor Sox9 has essential roles in successive steps of the chondrocyte differentiation pathway and is required for expression of Sox5 and Sox6. Genes Dev 16, 2813–2828 (2002).

59. Rafipay, A., Berg, A. L. R., Erskine, L. & Vargesson, N. Expression analysis of limb element markers during mouse embryonic development. Developmental Dynamics 247, 1217–1226 (2018).

60. Montero, J. A. et al. Sox9 Expression in Amniotes: Species-Specific Differences in the Formation of Digits. Front Cell Dev Biol 5, 23 (2017).

61. Luria, V., Krawchuk, D., Jessell, T. M., Laufer, E. & Kania, A. Specification of Motor Axon Trajectory by Ephrin-B:EphB Signaling: Symmetrical Control of Axonal Patterning in the Developing Limb. Neuron 60, 1039–1053 (2008).

62. Hutcheson, D. A., Zhao, J., Merrell, A., Haldar, M. & Kardon, G. Embryonic and fetal limb myogenic cells are derived from developmentally distinct progenitors and have different requirements for β-catenin. Genes Dev. 23, 997–1013 (2009).

63. Kardon, G., Campbell, J. K. & Tabin, C. J. Local Extrinsic Signals Determine Muscle and Endothelial Cell Fate and Patterning in the Vertebrate Limb. Developmental Cell 3, 533–545 (2002).

64. Sassoon, D. et al. Expression of two myogenic regulatory factors myogenin and MyoDl during mouse embryogenesis. Nature 341, 303–307 (1989).

65. Baumgartner, M. et al. Minor spliceosome inactivation causes microcephaly, owing to cell cycle defects and death of self-amplifying radial glial cells. Development 145, dev166322 (2018).

66. Beauchamp, M.-C., Alam, S. S., Kumar, S. & Jerome-Majewska, L. A. Spliceosomopathies and neurocristopathies: Two sides of the same coin? Developmental Dynamics 249, 924–945 (2020).

67. Griffin, C. & Saint-Jeannet, J.-P. Spliceosomopathies: Diseases and mechanisms. Dev Dyn 249, 1038–1046 (2020).

68. Chen, H. & Johnson, R. L. Dorsoventral patterning of the vertebrate limb: a process governed by multiple events. Cell Tissue Res 296, 67–73 (1999).

69. White, A. K. et al. Trp53 ablation fails to prevent microcephaly in mouse pallium with impaired minor intron splicing. Development 148, dev199591 (2021).

70. Kim, D., Langmead, B. & Salzberg, S. L. HISAT: a fast spliced aligner with low memory requirements. Nat Methods 12, 357–360 (2015).

71. Metsalu, T. & Vilo, J. ClustVis: a web tool for visualizing clustering of multivariate data using Principal Component Analysis and heatmap. Nucleic Acids Res 43, W566–570 (2015).

72. Cox, J. & Mann, M. MaxQuant enables high peptide identification rates, individualized p.p.b.-range mass accuracies and proteome-wide protein quantification. Nat Biotechnol 26, 1367–1372 (2008).

73. Hao, Y. et al. Integrated analysis of multimodal single-cell data. Cell 184, 3573–3587.e29 (2021).

74. Fiala, J. C. Reconstruct: a free editor for serial section microscopy. J Microsc 218, 52–61 (2005).

