## Supplementary Figures and Tables for "Integrated multi-omics reveals minor spliceosome inhibition causes molecular stalling and developmental delay of the mouse forelimb"

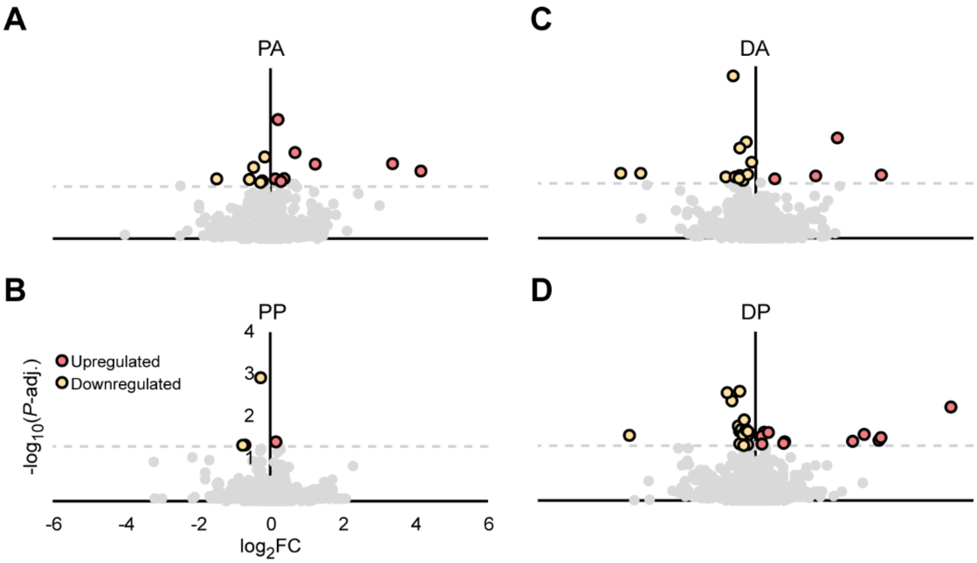


**Supplementary Figure 1. Differential protein expression in mutant microdomains.** (A-D) Volcano plots showing differentially expressed proteins in the mutant PA (A), PP (B), DA (C), and DP (D) microdomains. Dashed line shows cut-off for significance. PA=proximal anterior; PP=proximal posterior; DA=distal anterior; DP=distal posterior; adj.=Benjamini-Hochberg adjusted; FC=fold-change.

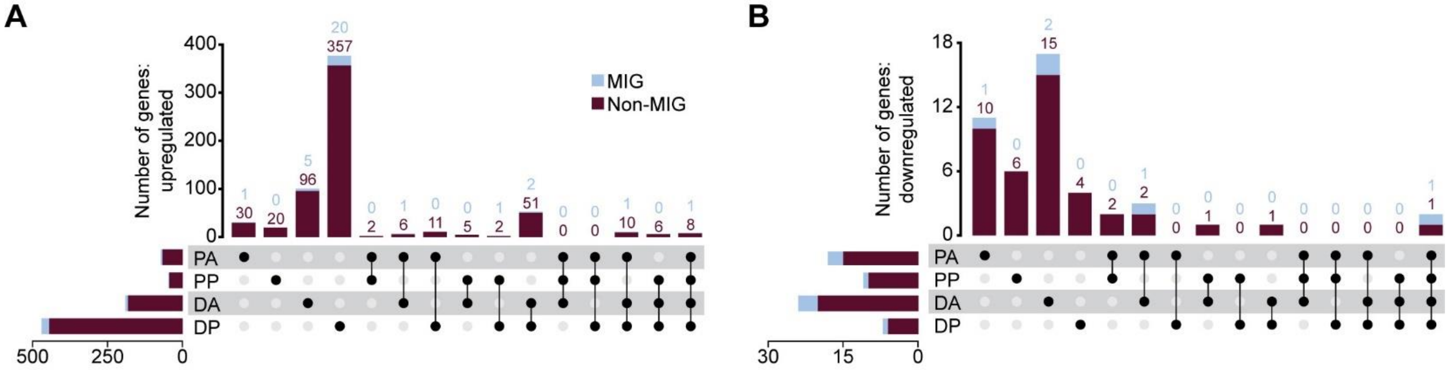


**C**

**Upregulated genes**

| Term (individual analysis) | Bin | No. genes | BH |
|---|---|---|---|
| KEGG: Prion disease | PA | 8 | 9.96E-03 |
| KEGG: Diabetic cardiomyopathy | PA | 7 | 1.12E-02 |
| KEGG: p53 signaling pathway | DA | 8 | 4.77E-04 |
| GO-CC: Cell surface | DA | 17 | 4.93E-02 |
| GO-MF: Protein binding | DP | 164 | 1.62E-05 |
| KEGG: p53 signaling pathway | DP | 11 | 8.70E-04 |
| GO-CC: Membrane | DP | 184 | 2.08E-03 |
| GO-CC: Collagen trimer | DP | 10 | 1.02E-02 |
| GO-BP: Negative regulation of cell growth | DP | 14 | 1.11E-02 |

**D**

**Upregulated genes**

| Term (overlap analysis) | Bin | No. genes | BH |
|---|---|---|---|
| KEGG: Prion disease | PA only | 5 | 3.16E-02 |
| KEGG: Oxidative phosphorylation | PA only | 4 | 3.16E-02 |
| GO-MF: Protein binding | DP only | 136 | 9.05E-05 |
| GO-CC: Membrane | DP only | 155 | 7.82E-04 |
| GO-MF: Nucleotide binding | DP only | 53 | 1.27E-02 |
| GO-MF: ATP binding | DP only | 48 | 1.46E-02 |
| GO-CC: Cytoplasm | DP only | 157 | 2.47E-02 |
| GO-MF: Extracellular matrix structural constituent | DP only | 10 | 3.58E-02 |
| KEGG: p53 signaling pathway | DA, DP only | 4 | 5.96E-02* |

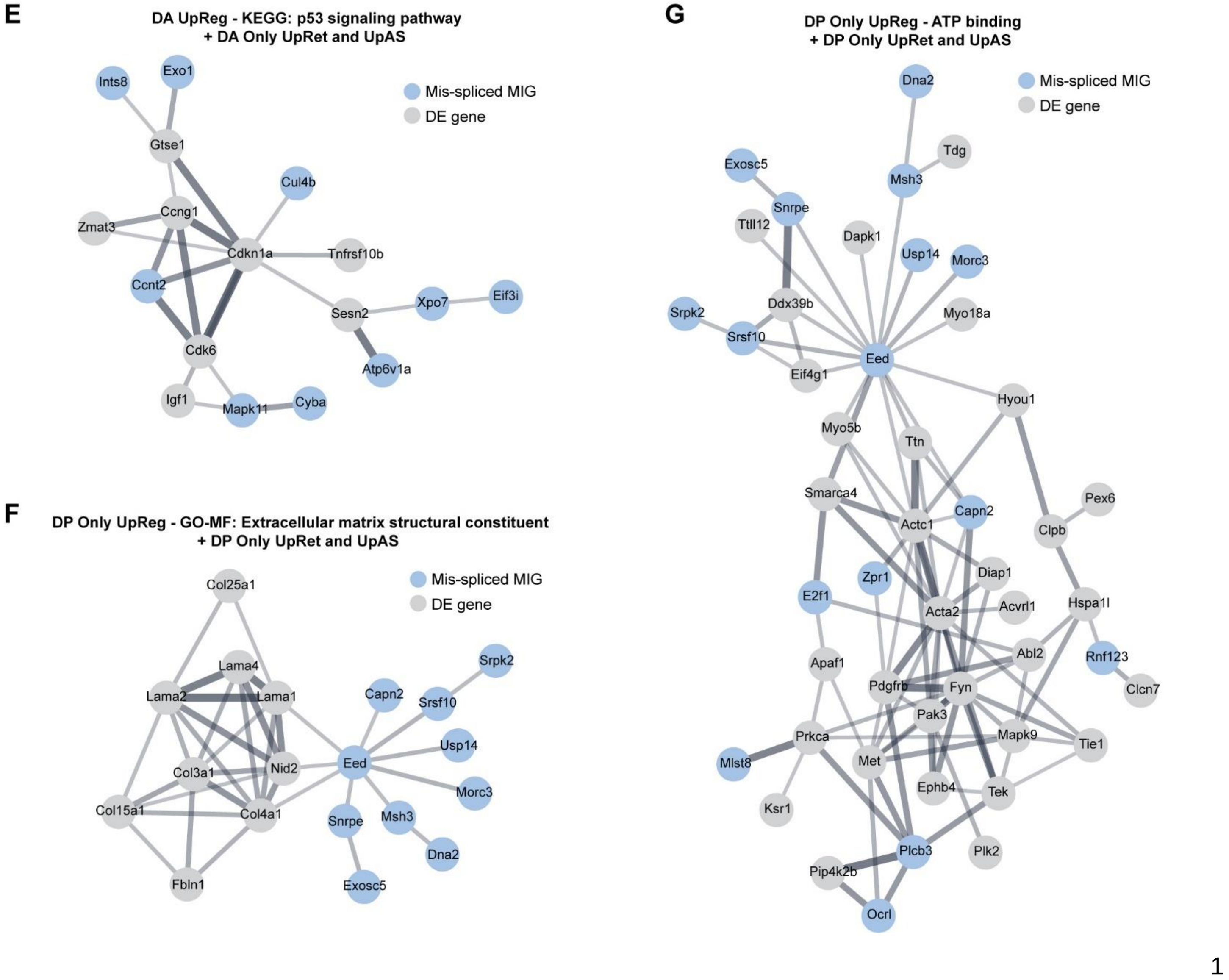

**Supplementary Figure 2. Spatially distinct gene expression changes interact directly with spatially mis-spliced MIGs.** (A-B) Upset plot showing number of upregulated (A) and downregulated (B) genes in each mutant microdomain when compared to its appropriate WT counterpart. Total set size located left of each sample. (C-D) Significantly enriched KEGG pathways and GO terms for genes upregulated in each head-to-head microdomain comparison (C) and after overlap analysis (D). *=slightly above significance threshold. (E) STRING network from genes upregulated in mutant DA microdomain that enrich for KEGG p53 signaling pathway along with MIGs showing upregulated retention (UpRet) or AS (UpAS) only in the DA microdomain, as identified in Fig. 2F, J. (F-G) STRING networks from genes upregulated only in mutant DP microdomain that enrich for extracellular matrix structural constituent (F) or ATP binding (G) submitted with MIGs showing UpRet or UpAS only in mutant DP microdomain as identified in Fig. 2F, J. Supplemental information for data shown in this figure can be found in Data File 4.

**A**

**WT**

| Domain | GOTerm | # Genes | BH-adj. *P* |
|---|---|---|---|
| Dis. | Proximal/distal pattern formation | 5 | 6.79E-8 |
| Dis. | Positive regulation of cell proliferation | 9 | 3.12E-2 |
| DA | Sequence specific DNA binding | 4 | 1.83E-2 |
| PA | Embryonic skeletal system morphogenesis | 5 | 8.74E-4 |
| PA | Extracellular region | 13 | 1.76E-3 |
| Prox. | Skeletal system development | 11 | 2.83E-4 |
| Prox. | Regulation of cell migration | 7 | 2.14E-2 |
| Prox. | Nervous system development | 14 | 2.84E-2 |
| Prox. | Skeletal muscle tissue development | 6 | 3.18E-2 |
| PP | Proteinaceous extracellular matrix | 5 | 1.52E-2 |
| Post. | Transcription regulatory region DNA binding | 3 | 4.44E-3 |
| DP | Pattern specification process | 5 | 8.64E-7 |
| DP | Hedgehog receptor activity | 2 | 1.39E-2 |

**B**

**Mut**

| Domain | GOTerm | # Genes | BH-adj. *P* |
|---|---|---|---|
| Dis. | Proximal/distal pattern formation | 9 | 3.35E-10 |
| Dis. | Negative regulation of canonical Wnt sig. | 6 | 6.90E-3 |
| PA | Embryonic skeletal system morphogenesis | 4 | 4.56E-4 |
| PA | Anterior/posterior pattern specification | 3 | 3.79E-2 |
| Prox. | Skeletal system development | 7 | 6.88E-3 |
| Prox. | Negative regulation of androgen receptor sig. | 4 | 8.07E-3 |
| Prox. | Neuron migration | 6 | 4.97E-2 |
| DP | Pattern specification process | 5 | 8.64E-7 |
| DP | Hedgehog receptor activity | 2 | 1.39E-2 |
| DP | Positive regulation of transcription | 4 | 3.53E-2 |

**Supplementary Figure 3. Biological pathway enrichment for genes with spatially restricted expression.** (A-B) DAVID gene ontology (GO) terms generated from spatially restricted genes for each domain in the WT (A) and mutant (B) *spatialPlot*. Sig.=signaling; Prox=proximal; Dis=distal; Ant=anterior; Post=posterior; PA=proximal anterior; PP=proximal posterior; DA=distal anterior; DP=distal posterior; BH-adj.=Benjamini-Hochberg-adjusted. Supplemental information for data shown in this figure can be found in Data File 5.

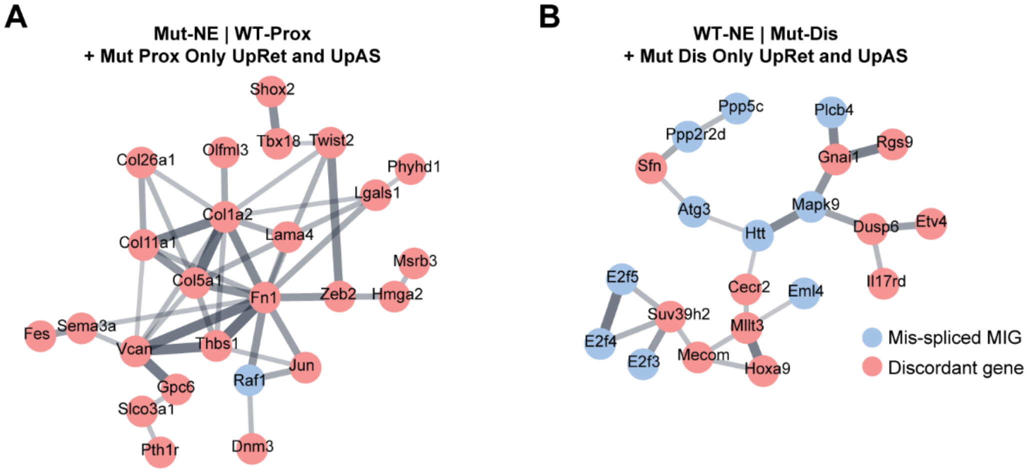


**Supplementary Figure 4. Spatially discordant genes form putative networks with spatially mis-spliced MIGs.** (A) STRING network showing connectivity of genes not enriched in the mutant transcriptome (Mut-NE) although proximally enriched in the WT transcriptome (WT-Prox) as shown in Fig. 3I with MIGs found to have upregulated retention (UpRet) or alternative splicing (UpAS) in proximal mutant microdomains (PA+PP) as shown in Fig. 2F, J. (B) STRING network showing connectivity of genes lacking spatial enrichment in the WT transcriptome (WT-NE) although distally enriched in the mutant transcriptome (Mut-Dis) as shown in Fig. 3K submitted along with MIGs found to have UpRet or UpAS in distal mutant microdomains (DA+DP) as shown in Fig. 2F, J.

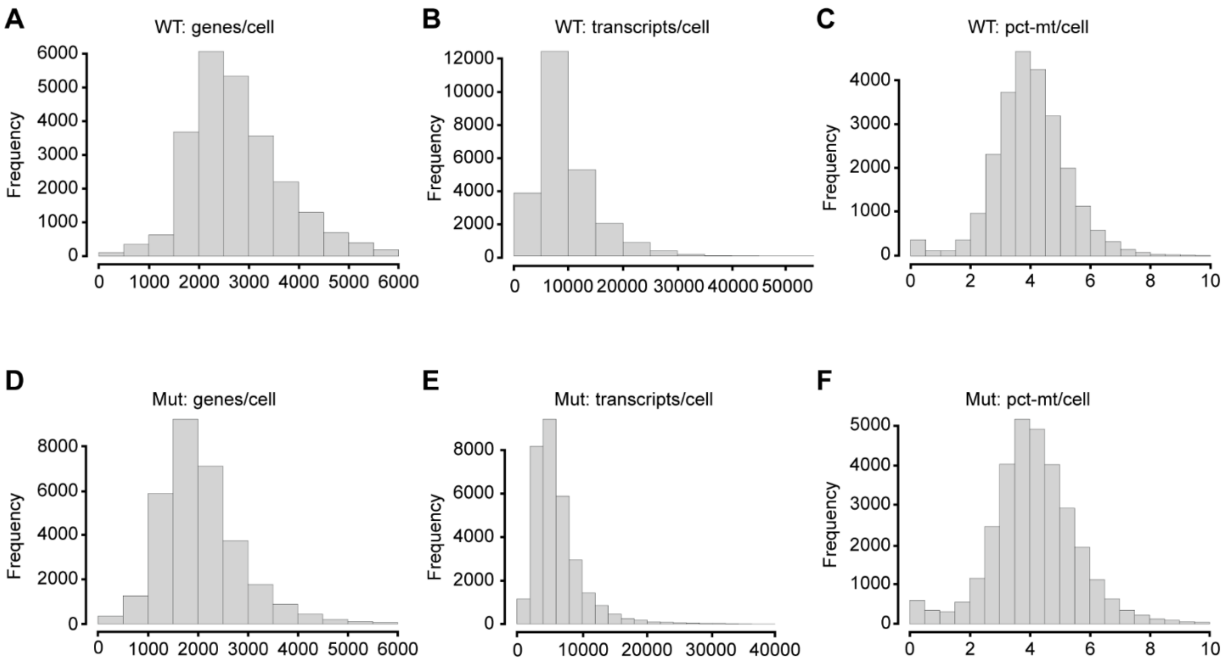


**Supplementary Figure 5. Single cell RNAseq data quality.** (A-E) Histograms showing number of genes per cell, transcripts per cell, and percent mitochondrial reads (pct-mt) per cell post-filtering for the WT (A-C) and mutant (D-F) datasets.

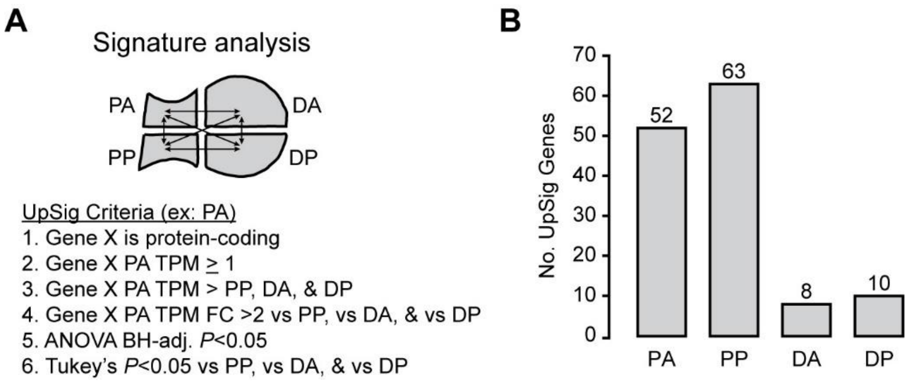


**Supplementary Figure 6. Obtaining microdomain RNAseq signature genes for addModuleScore.** (A) Criteria used for signature analysis with microdomain RNAseq. (B) Total number of signature genes identified for each WT microdomain; only the top eight genes were used for module scoring. Supplemental information for data shown in this figure can be found in Data File 6.

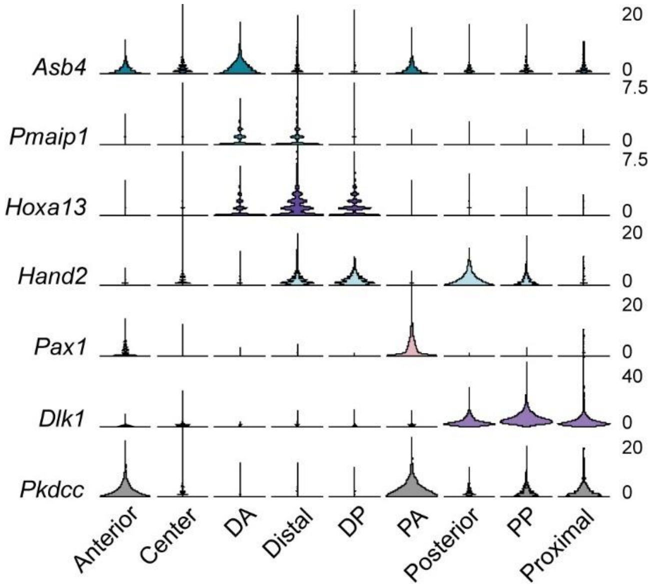


**Supplementary Figure 7. Validating single cell spatial binning.** Violin plots showing expression of known regional- or domain-specific spatial markers for each of the spatial single cell bins.

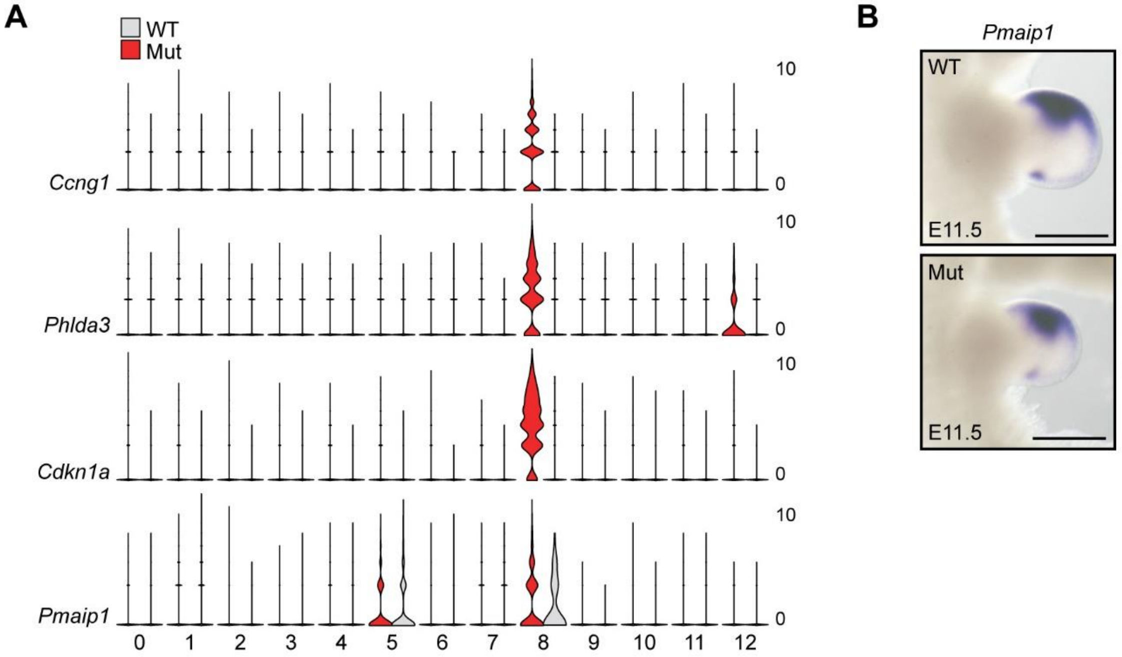


**Supplementary Figure 8. Single cell RNAseq reveals cluster of cells undergoing P53-mediated apoptosis in mutant forelimb.** (A) Violin plot for top transcriptional markers of cluster 8. (B) WISH for *Pmaip1* in E11.5 WT and mutant (Mut) forelimb. Scale bar shows 100 µm.

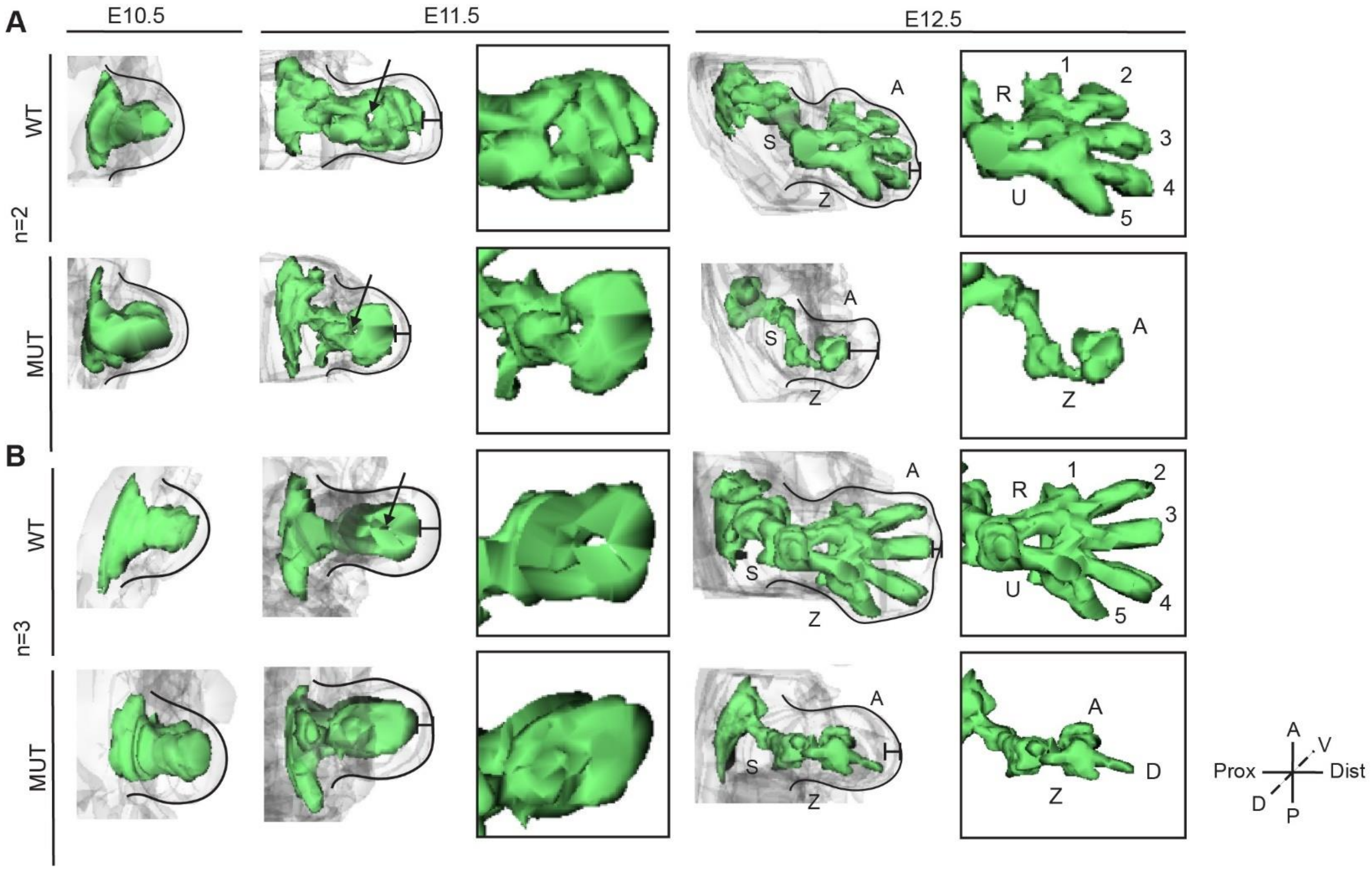


**Supplementary Figure 9. Aberrant chondrogenesis in the U11-null forelimb.** (A-B) Three-dimensionally reconstructed immunofluorescence staining pattern for Sox9 in N=2 (A) and N=3 (B) WT and mutant (Mut) forelimbs at E10.5, E11.5, and E12.5. Arrowheads show zeugopod bifurcation. Reconstructions are scaled between equivalent WT and Mut pairs, but not between time points or N-values. D=dorsal, V=ventral, An=anterior, Po=posterior, S=stylopod, Z=zeugopod, A=autopod, R=radius, U=ulna, 1-5=digits, D=digit.

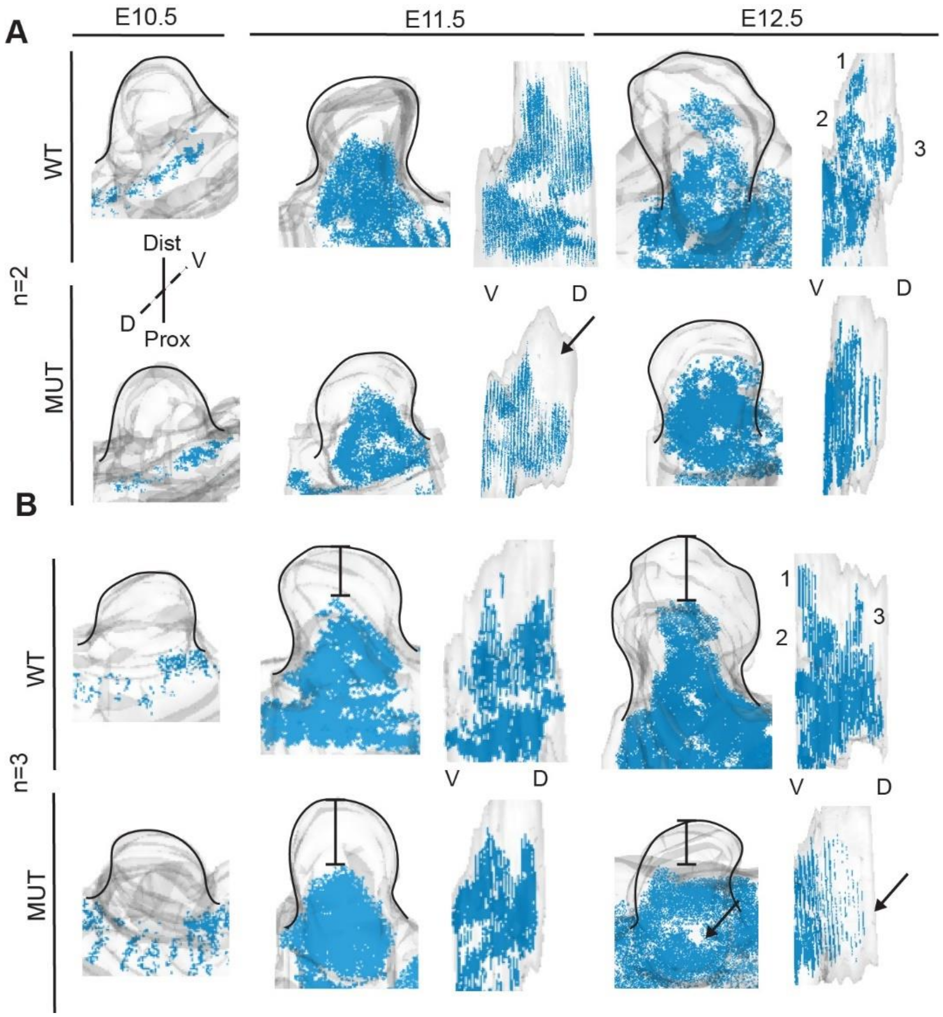


**Supplementary Figure 10. Aberrant myogenesis in the U11-null forelimb.** (A-B) Three-dimensionally reconstructed immunofluorescence staining pattern for Myod1 in N=2 (A) and N=3 (B) WT and mutant (Mut) forelimbs at E10.5, E11.5, and E12.5. Arrowhead shows lack of dorsal muscle mass in Mut. Reconstructions are scaled between equivalent WT and Mut pairs, but not between time points or N-values. Dist=distal, Prox=proximal, D=dorsal, V=ventral, 1-3=muscle masses.

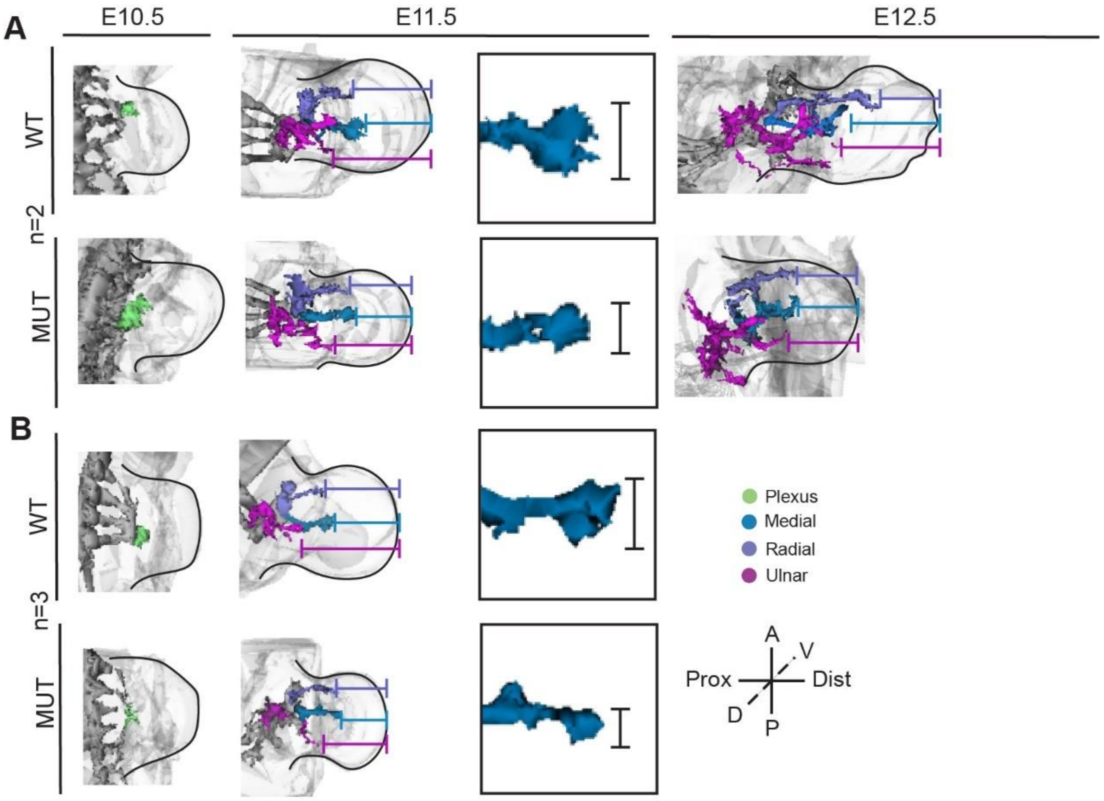


**Supplementary Figure 11. Aberrant axonogenesis in the U11-null forelimb.** (A-B) Three-dimensionally reconstructed immunofluorescence staining pattern for Neurofilament-M in N=2 (A) and N=3 (B) WT and mutant (Mut) forelimbs at E10.5, E11.5, and E12.5. Reconstructions are scaled between equivalent WT and Mut pairs, but not between time points or N-values. Prox=proximal, Dist=distal, Ant=anterior, Pos=posterior, D=dorsal, V=ventral.

**Table 1. Primer sequences used for RT-PCR.**

| Primer | Sequence (5' to 3') |
|---|---|
| *Fam92a* Forward | CGAAATCAACCTGTATGCCTCTACCGA |
| *Fam92a* Reverse | TGCGGGTAGCATCAATCGTGGCT |
| *Hnrnpm* Forward | GGCTGGAAGACTTGGAAGCACAGTAT |
| *Hnrnpm* Reverse | GTCTCCCTTTGGTAAAGCCCTCTC |
| *Exosc1* Forward | CTCTAGCATCAACTCACGGTTTGCCA |
| *Exosc1* Reverse | CTCGTTTTCAGCAGTAGTCAGCAGG |
| *Cdk5* Forward | GGCAATGATGTGGATGACCAGCTGA |
| *Cdk5* Reverse | CACCAAGGATGTTGTAGCTGGGTACAT |
